# *Smchd1* is a maternal effect gene required for autosomal imprinting

**DOI:** 10.1101/2020.01.20.913376

**Authors:** Iromi Wanigasuriya, Quentin Gouil, Sarah A. Kinkel, Andrés Tapia del Fierro, Tamara Beck, Ellise E.A. Roper, Kelsey Breslin, Jessica Stringer, Karla Hutt, Heather J. Lee, Andrew Keniry, Matthew E. Ritchie, Marnie E. Blewitt

**Affiliations:** Walter and Eliza Hall Institute of Medical Research, Parkville, Australia; The Department of Medical Biology, The University of Melbourne, Parkville, Australia; Faculty of Health and Medicine, The University of Newcastle, Australia; Monash Biomedicine Discovery institute, Monash University, Clayton, Australia; The Department of Mathematics and Statistics, The University of Melbourne, Parkville, Australia

**Keywords:** Smchd1, maternal effect gene, genomic imprinting, germline methylation imprints, allele-specific expression, imprinted H3K27me3

## Abstract

Genomic imprinting establishes parental allele-biased expression of a suite of mammalian genes based on parent-of-origin specific epigenetic marks. These marks are under the control of maternal effect proteins supplied in the oocyte. Here we report the epigenetic repressor *Smchd1* as a novel maternal effect gene that regulates imprinted expression of 16 genes. Most Smchd1-sensitive genes only show loss of imprinting post-implantation, indicating maternal Smchd1’s long-lived epigenetic effect. Sm-chd1-sensitive genes include both those controlled by germline polycomb marks and germline DNA methylation imprints; however, Smchd1 differs to other maternal effect genes that regulate the latter group, as Smchd1 does not affect germline DNA methylation imprints. Instead, Smchd1-sensitive genes are united by their reliance on polycomb-mediated histone methylation marks as germline or secondary imprints. We propose that Smchd1 translates these imprints to establish a heritable chromatin state required for imprinted expression later in development, revealing a new mechanism for maternal effect genes.

## Introduction

Genomic imprinting describes the process that enables monoallelic expression of a set of genes according to their parent of origin (1, 2). Classically, these imprinted genes are located in clusters. Disruption of imprinting at specific gene clusters leads to imprinting disorders. For example, loss of imprinting at the *SNRPN* cluster is responsible for Prader Willi syndrome or Angelman syndrome, and loss of imprinting at the *IGF2*/*H19* and *KCNQ1* clusters is responsible for Beckwith-Wiedemann syndrome (3–10). Imprinted expression is enabled by germline DNA methylation or histone methylation imprints that resist preimplantation reprogramming and therefore retain their parent-of-origin specific marks (11–14). Evidence to date suggests that proteins found in the oocyte establish the imprints and enable them to resist preimplantation reprogramming. In this way these maternally derived proteins control imprinted expression. To date only a small number of such maternal effect genes have been identified, including *Trim28*, *Zfp57*, *Stella*, *Rlim*, *Nlrp2*, *Dnmt3L*, *Dnmt1o* and *Eed* (13–23).

Smchd1 is an epigenetic modifier, the zygotic form of which is required for both X chromosome inactivation (24, 25) and silencing of clustered autosomal loci, including genes at the *Snrpn* and *Igf2r-Airn* imprinted clusters (26–29). SMCHD1 function is also relevant in the context of disease (30). Heterozygous mutations in *SM-CHD1* are found in two distinct human disorders: loss of function mutations in facioscapulohumeral muscular dystrophy (FSHD) (31), and potentially gain of function mutations in the rare craniofacial disorder Bosma arhinia and microphthalmia (BAMS) (32–34). The role of SMCHD1 in normal development and disease has led to an increase in interest in how and when it contributes to gene silencing at each of its genomic targets.

Recent work by our group and others has shown that Smchd1 is required for long range chromatin interactions on the inactive X chromosome (28, 35, 36), and at its autosomal targets including the imprinted loci (28). In cells lacking Smchd1, other epigenetic regulators are enriched at Smchd1 targets, for example, CCCTC-binding factor (Ctcf) binding is enhanced at the inactive X chromosome and the clustered protocadherins (26, 35, 36). The inactive X also shows an accumulation of polycomb repressive complex 2-mediated histone 3 lysine 27 trimethylation (H3K27me3) in the absence of Smchd1 (28). These data led to a model where Smchd1-mediated chromatin interactions insulate the genome against the action of other epigenetic regulators, such as Ctcf (26, 28, 35, 36); however, it is not known whether this model holds true for all Smchd1 target genes.

Based on Smchd1’s known role at some imprinted clusters (26–29) and its expression in the oocyte (37), we sought to test the role of oocyte-supplied maternal Sm-chd1 in regulating imprinted gene expression.

Here we show that *Smchd1* is a novel maternal effect gene, required for imprinted expression of at least 16 genes. Unlike other known maternal effect genes, with the exception of *Eed*, deletion of maternal *Smchd1* does not disrupt germline DNA methylation imprints during preimplantation development or later in gestation. Interestingly, some genes that show perturbed imprinted expression later in development in the absence of maternal Smchd1 are not expressed in the preimplantation period.

These data lead us to propose that Smchd1 operates via a distinct mechanism for a maternal effect gene, where, based on the imprints, it establishes an epigenetic memory later required for imprinted expression.

## Results

### Genetic deletion of maternal *Smchd1*

To determine whether maternally derived Smchd1 is required for autosomal imprinting, we deleted *Smchd1* in the oocyte using MMTV-Cre or Zp3-Cre (Fig. 1a). Contrary to previous reports where Smchd1 depletion by siRNA knockdown in preimplantation embryos resulted in reduced blastocyst formation, hatching and survival to term (37, 38), we found no embryonic or preweaning lethality following maternal deletion of *Smchd1* (Fig. S1a).

**Fig. 1.**
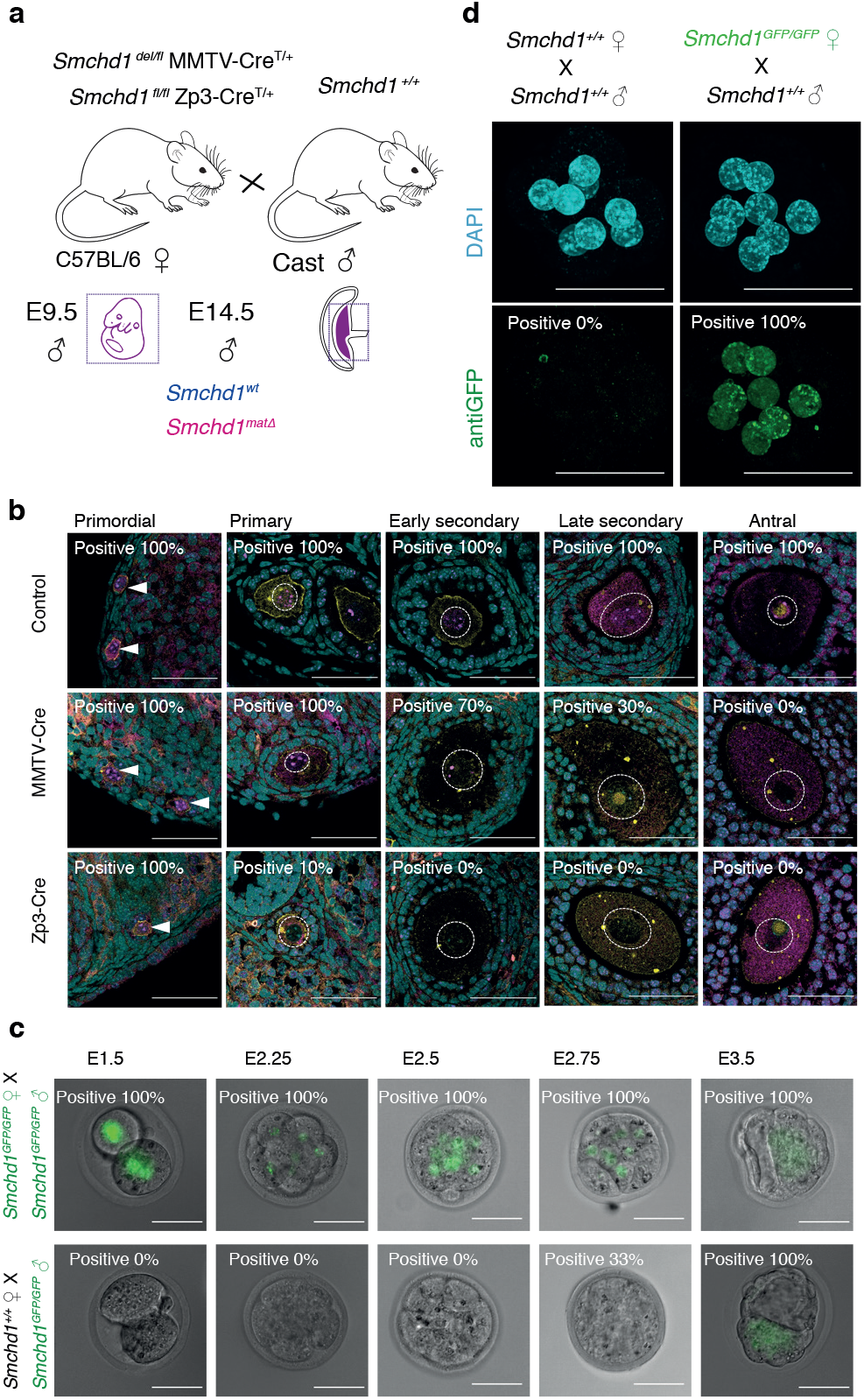
Maternal deletion of *Smchd1* during oocyte development depletes Smchd1 until the 16-cell embryonic stage. **a.** Schematic for maternal deletion of *Smchd1*. **b.** Deletion of *Smchd1* in oocyte development with MMTV-Cre and Zp3-Cre. Arrowheads indicate primordial follicle oocyte nuclei, white dotted lines surround primary–antral follicle oocytes. Smchd1 (magenta), c-Kit (yellow), DAPI (cyan). n = 15–27 sections for 2 ovaries per cohort. A total of 5–20 follicles were observed for primordial to late secondary stages and 2–3 antral follicles for each genotype. **c.** Detection of paternal Smchd1-GFP from day 1.5 to 3.5 in preimplantation embryos (n = 2–5). *Smchd1*^GFP/GFP^ embryos were used as positive controls. **d.** Detection of maternal Smchd1-GFP in the nuclei of E2.5 (8-cell) embryos, with *Smchd1*^+/+^ used as negative controls (n = 1–2). Nuclei marked with DAPI (cyan). Scale bar: 50 μm.

Using MMTV-Cre, Smchd1 was deleted progressively from the early secondary to antral follicle stages of oocyte development, whereas Zp3-Cre deleted earlier, in the primary follicle (Fig. 1b). Using our Smchd1-GFP fusion knock-in mouse (28), we found that paternally encoded native Smchd1-GFP protein was not detectable until the 16-cell stage (Fig. 1c). We confirmed the expression by immunofluorescence and found very low levels of paternal Smchd1 switching on from the 8-cell stage (Fig. S1b), but its expression is much lower than maternal Smchd1 at this time (Fig. 1d). Together these data suggest maternal Smchd1 is the primary source of protein until at least the 32-cell stage. These data define the period of maternal deletion of *Smchd1* in each model.

Using the same approach we also confirmed that maternal Smchd1 localises to the nucleus in the preimplantation period (Fig. 1d), as expected if Smchd1 plays a functional role in gene silencing at this time.

### Maternal Smchd1 regulates the imprinting of known Smchd1 targets

Using this system, we analysed the effect of maternal Smchd1 on expression using RNA-seq in male *Smchd1* maternally deleted (*Smchd1*^matΔ^) E9.5 embryos and the embryonic portion of E14.5 placentae. Females were not examined due to the confounding role of Smchd1 in X chromosome inactivation. We found only five consistently differentially expressed genes between *Smchd1*^matΔ^ and control placental samples from both Cre models (Fig. S1c-e), and limited differential expression in the E9.5 embryos analysed (MMTV-Cre, Fig. S1f). These data suggest removal of maternal Sm-chd1 does not have a striking effect on gene expression. These data are consistent with no observable effect on viability by deletion of maternal *Smchd1* (Fig. S1a).

To examine the role of maternal Smchd1 in imprinted expression we took advantage of the single nucleotide polymorphisms within the same C57BL/6 x CAST/EiJ F1 samples for allele-specific analyses (Fig. 1a). We observed consistent effects on imprinted expression with both Cre strains, suggesting that the stage of deletion during oogenesis does not influence outcome (Fig. S2a). Therefore, we pooled data from both Cre strains for statistical analyses. We imposed a 5% FDR as well as an absolute difference in the proportion of the RNA derived from the silenced allele greater than 5% to call differential imprinted expression.

E14.5 *Smchd1*^matΔ^ placentae displayed partial loss of imprinted expression at known Smchd1-sensitive genes in the *Snrpn* cluster (*Magel2*, *Peg12* and *Ndn*, Fig. 2a, Additional file 1) and the *Igf2r-Airn* cluster (*Slc22a3* and *Pde10a*, Fig. 2b, Additional file 1), shown as an increase in the proportion of the RNA derived from the silenced allele (Fig. S2b).

**Fig. 2.**
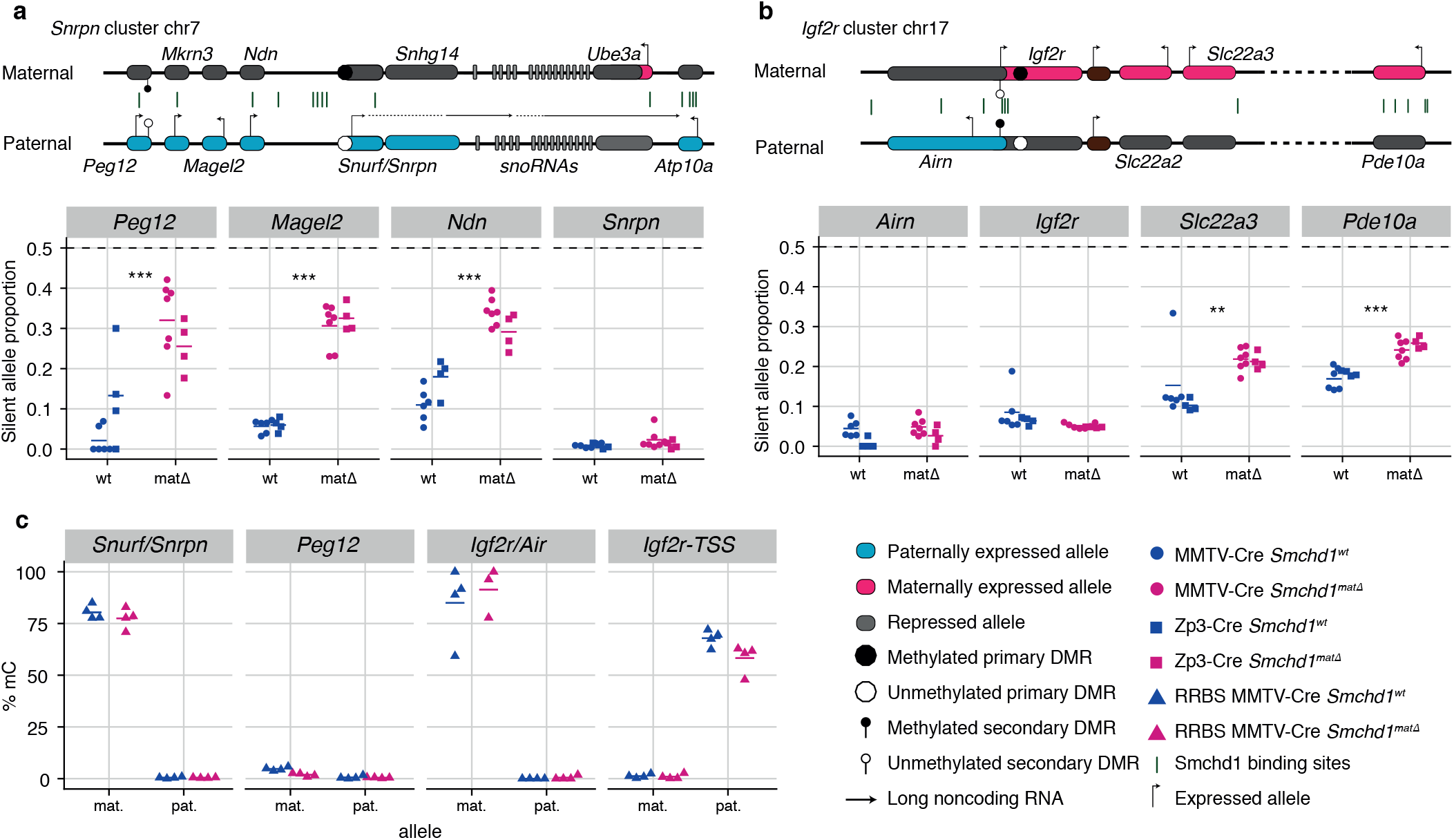
Maternal deletion of *Smchd1* results in partial loss of imprinting at known Smchd1-sensitive clusters. **a-b.** Expression of the silent allele as a proportion of total expression of the gene, obtained by allele-specific RNA-seq from the embryonic portion of the placenta of Smchd1 maternal null conceptuses. Green lines indicate Smchd1 binding sites determined by ChIP-seq. Expression data for **a.** *Snrpn* cluster genes on chromosome 7 and **b.** *Igf2r* cluster genes on chromosome 17. **c.** Percentage methylation (% mC) on the maternal and paternal alleles at primary and secondary DMRs at *Snrpn* and *Igf2r* clusters in Smchd1 maternal null (matΔ) and wild-type (wt) placental samples. * *p* < 0.05, ** *p* < 0.01, *** *p* < 0.001, when the difference in mean silent allele proportions between genotypes is of at least 5%. RNA-seq: n = 6 MMTV-Cre, n = 4 Zp3-Cre wild-type (wt); n = 7 MMTV-Cre, n = 4 Zp3-Cre Smchd1 maternal null (matΔ) E14.5 placental samples. RRBS n = 4 MMTV-Cre for both matΔ and wt E14.5 placental samples.

Far fewer genes are imprinted in the embryo compared with the placenta, however partial loss of imprinted expression was also observed at *Peg12* and *Ndn* in E9.5 embryos (Fig. S2c, Additional file 1).

### Maternal deletion of *Smchd1* does not affect germline DMRs

We analysed DNA methylation at imprinted differentially methylated regions (DMRs) by reduced representation bisulfite sequencing (RRBS) in male E14.5 placentae and found that despite partial loss of imprinted expression in *Smchd1*^matΔ^ samples, DNA methylation at germline DMRs was maintained (Fig. 2c, Additional file 2). We and others previously showed that *Smchd1* null embryos display hypomethylation of the *Peg12* secondary DMR (27, 29), however this region is not methylated in placenta so we could not directly compare with our current data.

These data show *Smchd1* is a maternal effect gene required for autosomal imprinting at genes where zygotic Smchd1 also plays a role, and this occurs without changes to germline DMR methylation.

### New imprinted genes under the control of maternal Smchd1

We next analysed all other imprinted genes and observed partial loss of imprinted expression of 8 additional genes across 6 chromosomal locations in placental samples. These included *Kcnq1* cluster genes *Tssc4* and *Ascl2* on chromosome 7, and *Jade1* and *Platr4* that are co-located on chromosome 3 (Fig. 3a and b, Additional file 1). Additionally, imprinted genes *Sfmbt2*, *Smoc1*, *Epop* and *Spp1* showed partial loss of imprinted expression (Fig. 3b and Fig. S3a, Additional file 1).

**Fig. 3.**
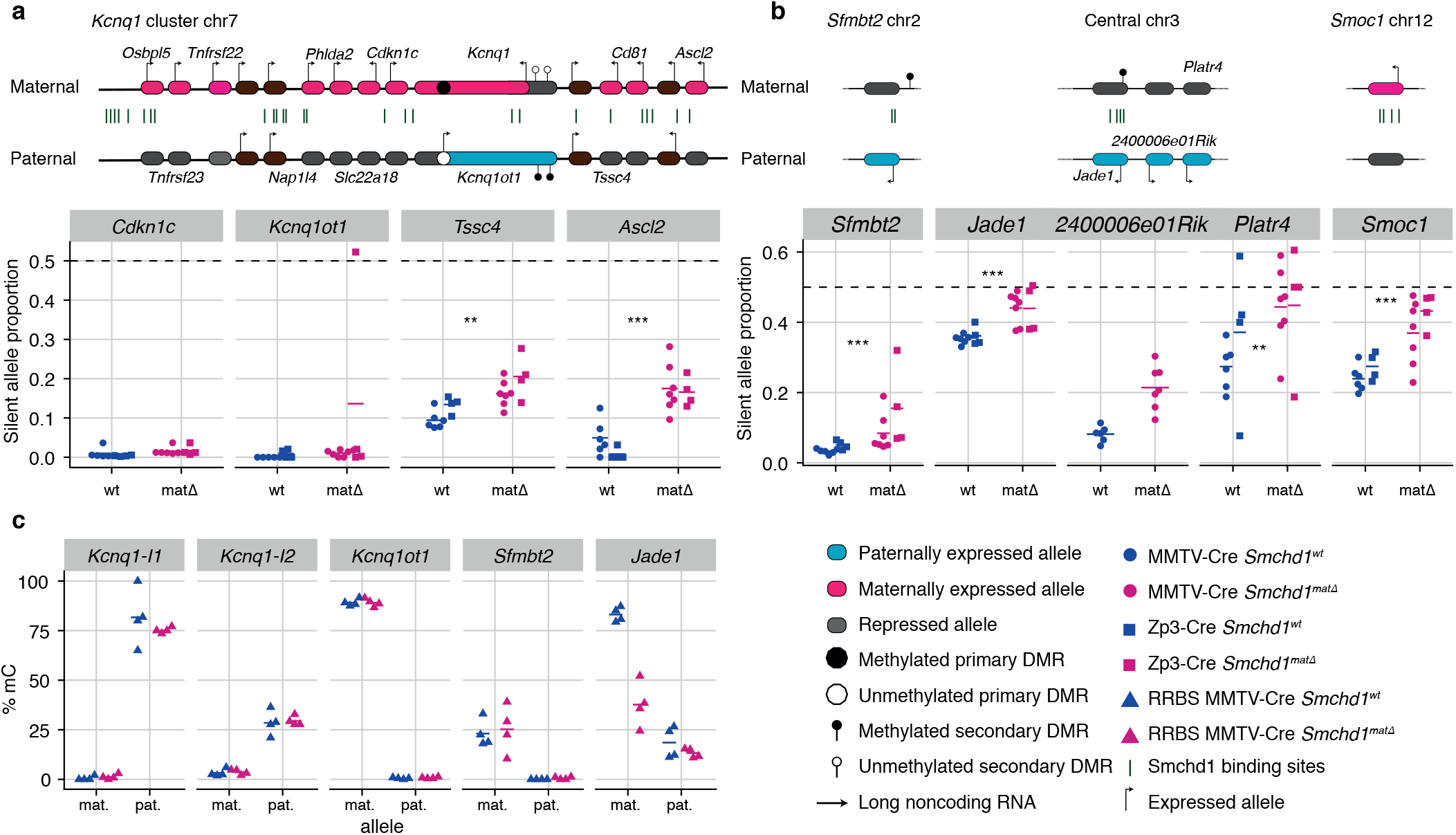
Maternal deletion of *Smchd1* results in loss of imprinting at the *Kcnq1* imprinted cluster and lone imprinted genes without changes to primary DMR methylation. **a-b**. Expression of the silent allele as a proportion of total expression of the gene, obtained by allele-specific RNA-seq from the embryonic portion of the placenta of *Smchd1* maternal null conceptuses. Green lines indicate Smchd1 binding sites determined by ChIP-Seq. Expression data for **a.** Kcnq1 cluster genes on chromosome 7, and **b.** at lone imprinted genes: *Sfmbt2*, *Jade1*, *2400006e01Rik* and *Platr4*, and *Smoc1*. **c.** Percentage methylation (% mC) for each parental allele at the DMRs for clusters and genes and samples shown in a. and b. *Kcnq1-I1*: *Kcnq1-Intergenic1*; *Kcnq1-I2*: *Kcnq1-Intergenic2*. * *p* < 0.05, ** *p* < 0.01, *** *p* < 0.001, when the difference in silent allele proportions is of at least 5%. RNA-seq: n = 6 MMTV-Cre, n = 4 Zp3-Cre wild-type (wt); n = 7 MMTV-Cre, n = 4 Zp3-Cre Smchd1 maternal null (matΔ) E14.5 placental samples. RRBS n = 4 MMTV-Cre for both matΔ and wt E14.5 placental samples.

Similarly to the *Snrpn* and *Igf2r-Airn* clusters, we detected no DMR hypomethylation, with one exception at the *Jade1* secondary DMR (Fig. 3c, Additional file 2). Supporting a role for Smchd1 in regulating these autosomal imprinted genes, we found Smchd1 binding sites across affected clusters and non-clustered imprinted genes in somatic cells (Fig. 2a-b, Fig. 3a-b, Fig. S3b). These data reveal a broader role for Smchd1 in autosomal imprinting than previously known.

### Both maternal and zygotic Smchd1 contribute to imprinted expression

Earlier work on *Zfp57* and *Trim28* showed that either maternal or zygotic deletions resulted in partially penetrant loss of imprinting, whereas deletion of both maternal and zygotic increased the penetrance, evidence that both the oocyte and zygotic supply of these proteins contribute to imprint regulation (15, 18). To understand the contribution of maternal versus zygotic Smchd1 at imprinted genes, we produced embryonic and placental samples that were wild-type, *Sm-chd1*^matΔ^, zygotic-deleted (*Smchd1*^zygΔ^) or maternal-and-zygotic-deleted for *Smchd1* (*Smchd1*^matzygΔ^). The consequence of our specific breeding scheme was that on average only half of the imprinted genes in each sample had the polymorphisms between parental alleles required for allele-specific analyses (Fig. S4a), and some genes had no informative samples.

Acknowledging this limitation, we did not identify any additional Smchd1-sensitive imprinted genes in *Sm-chd1*^zygΔ^ or *Smchd1*^matzygΔ^ samples compared with the maternal deletion alone (Additional file 1). In general, we observed a stronger effect of zygotic compared with maternal *Smchd1* deletion on imprinted expression (Fig. S4b-d). *Peg12*, *Magel2* and *Ndn* showed complete loss of imprinted expression in *Smchd1*^zygΔ^ and *Smchd1*^matzygΔ^ embryos and placentae, compared with only partial loss of imprinted expression in *Smchd1*^matΔ^ samples (Fig. 4a, and Fig. S4b). *Slc22a3*, *Pde10a* and *Sfmbt2* in *Smchd1*^zygΔ^ samples also showed significantly higher loss of imprinted expression compared to *Sm-chd1*^matΔ^ samples in placentae (Fig. 4b and c). These data suggest that maternal and zygotic Smchd1 play partly overlapping roles at imprinted genes.

**Fig. 4.**
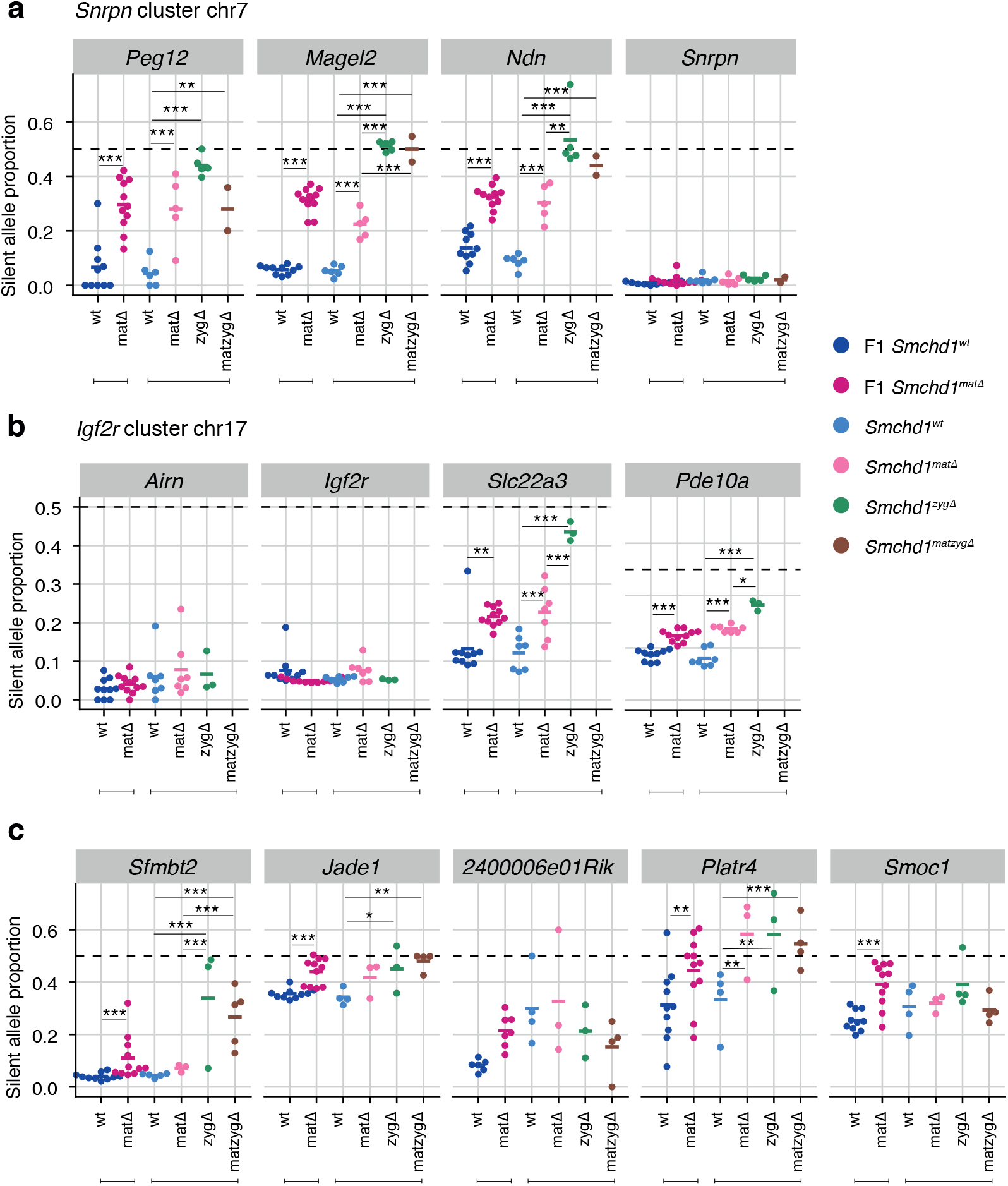
Zygotic deletion of *Smchd1* results in more severe loss of imprinting at genes sensitive to oocyte deletion of *Smchd1*. MMTV-Cre *Smchd1* oocyte-deleted data (F1 wt, F1 matΔ) from Figs.1 and 2, along with samples produced to compare *Smchd1* wild-type (wt), oocyte-deleted (matΔ), zygote-deleted (zygΔ), and oocyte-and-zygote-deleted (matzygΔ) genotypes. Samples from the embryonic portion of the placenta and expression of the silent allele is shown as a proportion of total expression of the gene, obtained by allele-specific RNA-seq. **a.** *Snrpn* cluster genes. **b.** *Igf2r-Airn* cluster genes. **c.** *Sfmbt2*, *Jade1*, *2400006e01Rik* and *Platr4*, and *Smoc1* genes. * *p* < 0.05, ** *p* < 0.01, *** *p* < 0.001, when the difference in silent allele proportions is at least 5%. RNA-seq for the zygotic deletion samples: n = 13 wt, n = 7 matΔ, n = 8 zygΔ, n = 6 matzygΔ for E14.5 MMTV-Cre placentae.

### Maternal Smchd1 sets up imprinted expression preimplantation without affecting germline DMRs nor requiring expression of target genes

To directly test the role of maternal Smchd1 in the preimplantation period, we carried out allele-specific methylome and transcriptome sequencing of *Smchd1*^matΔ^ and control 16-cell stage embryos, a period when maternally derived Smchd1 provides the primary supply of Smchd1 protein (Fig. 1c). Although we found minimal differential expression genome-wide (Fig. S5a), we found 89 genes with imprinted expression at this stage (Additional file 1), 4 of which showed significant loss of imprinted expression in the *Smchd1*^matΔ^ samples (Fig. 5a and b, Fig. S5b and c).

**Fig. 5.**
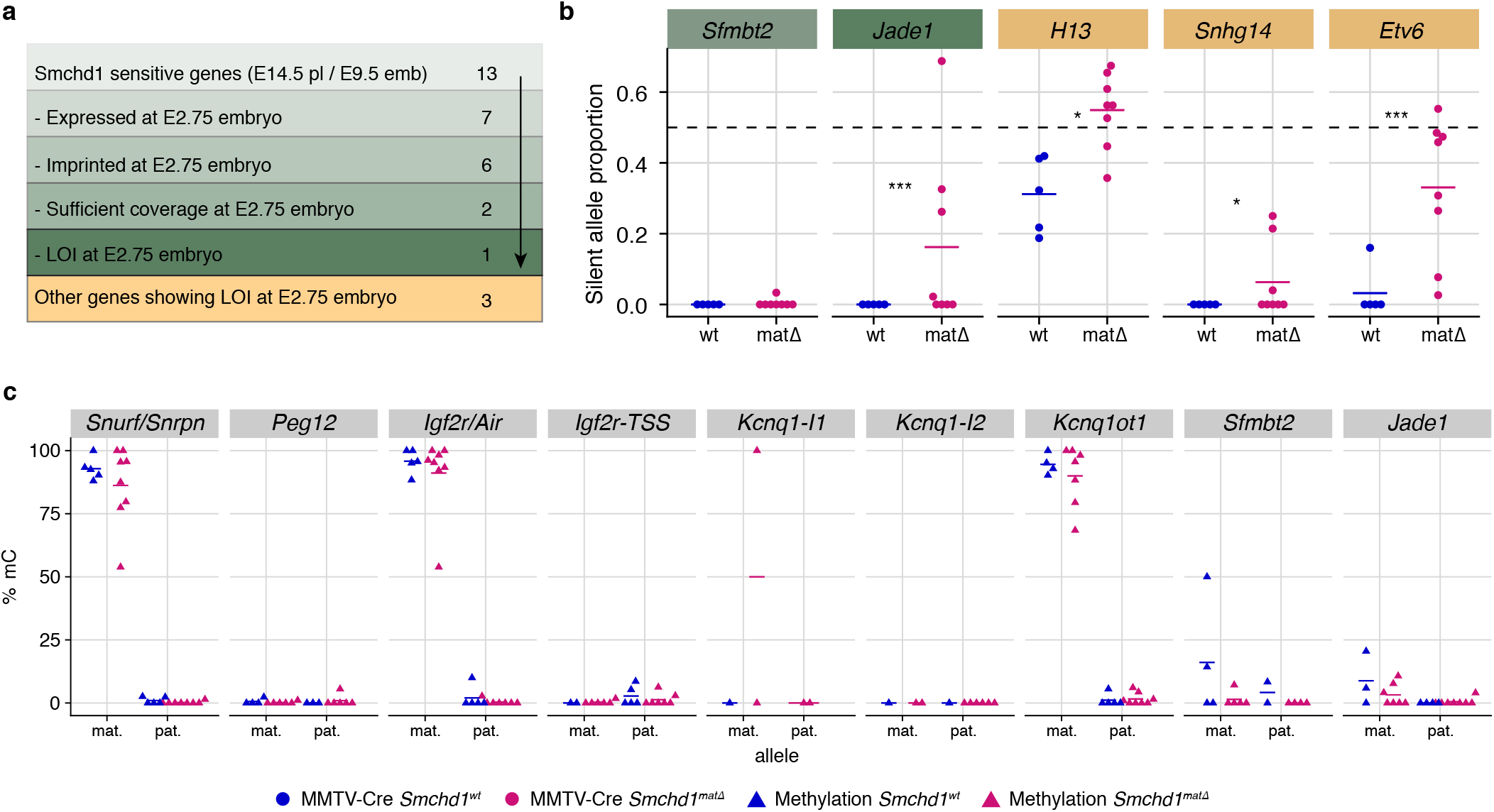
Maternal Smchd1 establishes an epigenetic memory required for imprinted gene expression. **a.** Summarised analysis of Smchd1-sensitive imprinted genes from E14.5 placentae and E9.5 embryos in E2.75-embryo transcriptome sequencing. **b.** Expression of the silent allele as a proportion of total expression of the gene, obtained by allele-specific RNA-seq from whole 2.75-day *Smchd1* maternal null (matΔ) and wild-type (wt) embryos. **c.**Percentage methylation (allele for the DMRs of Smchd1-sensitive imprinted clusters and genes in *Smchd1* maternal null (matΔ) compared with control (wt E2.75 embryos. *Kcnq1-I1*: *Kcnq1-Intergenic1*; *Kcnq1-I2*: *Kcnq1-Intergenic2*. * *p* < 0.05, ** *p* < 0.01, *** *p* < 0.001, when the difference in silent allele proportions is at least 5%. n = 5 wt and n = 8 matΔ E2.75 embryos.

Only two of the 13 maternal Smchd1-sensitive imprinted genes identified later in gestation were able to be analysed for differential imprinted expression at this time based on both expression and coverage of informative SNPs. One of these two genes, *Jade1*, showed loss of imprinted expression, while the other, *Sfmbt2*, did not yet show an effect.

These data suggest Smchd1 is already necessary for imprinting of *Jade1*, but instead is required for maintenance of imprinted expression for *Sfmbt2* (Fig. 5a and b). Since seven maternal Smchd1-sensitive genes were not yet expressed or imprinted, these data suggest that maternal Smchd1 imparts a long-lasting effect, independent of imprinted expression in the preimplantation period.

We analysed DNA methylation in the 16-cell embryos and again found no differences at germline DMRs (Fig. 5c, Additional file 2). This is surprising given that all other known maternal effect genes that regulate the same clusters of imprinted genes do so via regulation of the germline DNA methylation imprints. However, no change in germline DMR methylation is consistent with the lack of Smchd1 binding at the germline DMRs in somatic cells (26, 28) (Fig. S3) and the observed loss of imprinting only occurring at selected genes within the *Snrpn*, *Kcnq1* and *Igf2r-Airn* clusters. These data suggest that maternal Smchd1 regulates imprinting via a mechanism that is distinct from that employed by classic maternal effect genes.

## Discussion

Here, we identified *Smchd1* as a novel maternal effect gene. For the first time we revealed new clustered and non-clustered imprinted genes that are sensitive to deletion of maternal *Smchd1*. We revealed that each of these genes are also regulated by zygotic Smchd1. Together these data dramatically expand the number of imprinted loci that are sensitive to Smchd1 removal.

Known maternal effect genes act to establish and maintain the germline imprints during the preimplantation period. Interestingly, our data suggest that maternal Smchd1 has a different mechanism of action, as although we observe loss of imprinted expression at three clusters controlled by germline DNA methylation imprints (*Snrpn*, *Kcnq1* and *Igf2r-Airn* clusters), we see no change in DNA methylation at these regions, either preimplantation or later in development. These data suggest that maternal Smchd1 operates via a new mechanism compared with the other known maternal effect genes.

The Smchd1-sensitive imprinted genes fall into two classes: those in clusters controlled by DNA methylation at the germline DMR (canonical imprinted genes), and more recently discovered imprinted genes regulated instead by H3K27me3 imprints (non-canonical imprinted genes).

For some non-canonical imprinted genes (*Smoc1*, *Jade1*, *Sfmbt2*), it has recently been shown that the H3K27me3 imprint is lost during preimplantation development and through an unknown mechanism the imprint is transferred into a secondary DNA methylation imprint (39). By contrast, the *Kcnq1* cluster, the *Igf2r-Airn* cluster and the *Magel2* region of the *Snrpn* cluster gain H3K27me3 in post-implantation tissues, dependent on the germline DMR imprint (39), and for the *Kcnq1* and *Igf2r-Airn* clusters through the action of imprinted long noncoding RNAs (40–43). Interestingly, *Dnmt1* knock-out studies have shown that while imprinting of genes located centrally in the *Kcnq1* cluster is maintained via germline DMR methylation, genes affected by maternal deletion of *Smchd1*, *Tssc4* and *Ascl2*, are maintained via histone modifications (44–46).

Therefore, what appears to unite the Smchd1-sensitive imprinted genes is their reliance on H3K27me3 either as a germline or secondary imprint. Based on two lines of evidence, we suggest that Smchd1 acts downstream of these H3K27me3 imprints.

Firstly, we find the most striking loss of imprinted expression at both non-canonical and canonical imprinted genes in *Smchd1*^zygΔ^ samples (Fig. 4). These samples likely have no disruption to Smchd1 levels in the oocyte or preimplantation and therefore the non-canonical H3K27me3 imprints should remain.

Secondly, we have previously found no effect on H3K27me3 in *Smchd1* zygotic null samples at the Sm-chd1-sensitive clustered imprinted genes, i.e. secondary H3K27me3 imprints (26). Together these data suggest Smchd1 acts downstream of H3K27me3 in mediating imprinted expression (Fig. 6). We have previously shown that for the inactive X chromosome, Smchd1 is recruited dependent on polycomb repressive complex 1 (47). Potentially, a related pathway exists at these imprinted clusters.

**Fig. 6.**
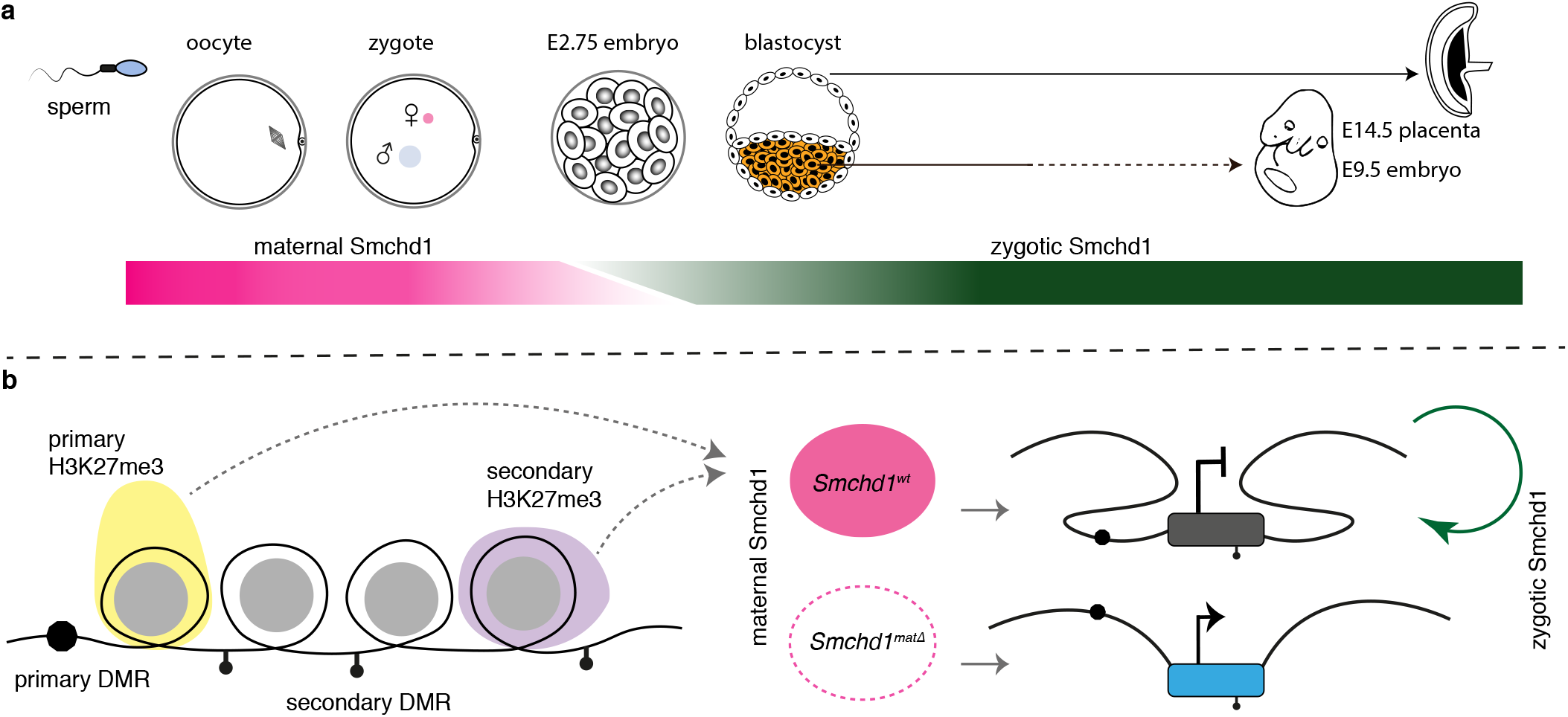
Smchd1 translates the imprints to establish a heritable chromatin state required for imprinted expression later in development. **a.** Developmental windows of activity of maternal and zygotic Smchd1. **b.** Proposed model illustrating the regulation of imprinted genes by Smchd1. Both oocyte and zygotic Smchd1 contribute to an epigenetic memory downstream of polycomb repressive histone marks.

We propose that Smchd1 plays a role in protecting the repressed imprinted genes from inappropriate activation. We and others have previously found that cells without Smchd1 have increased Ctcf binding (26, 35, 36). At Sm-chd1-sensitive imprinted genes we have found increased H3K4me2 or H3K4me3 (26, 29). Therefore, we suggest that Smchd1 may ensure that H3K27me3 imprints result in transcriptional silencing by preventing the action of Ctcf or other epigenetic activators via an insulating mechanism.

## Conclusions

We have discovered *Smchd1* is a novel maternal effect gene required for genomic imprinting. Recently, we and others discovered that zygotic Smchd1 mediates long-range chromatin interactions at its target genes, both imprinted regions and elsewhere in the genome (28, 35, 36). Based on the data presented here, we propose that Smchd1 is required to translate germline imprints into a chromatin state required to silence expression at select imprinted clusters. Given that many Smchd1-sensitive imprinted genes are not yet expressed in the preimplantation period, we hypothesize the chromatin state that maternal Smchd1 establishes, provides a long-lived epigenetic memory (Fig. 6). This work opens a new avenue to understand how imprinted expression is established during development and may be relevant for patients with SMCHD1 mutations.

## Methods

### Mouse strains and genotyping

The MMTV-Cre transgene line A (48) was backcrossed for more than 10 generations onto the C57BL/6 background from the FVB/N background for use in this study. This was used in combination with a *Smchd1* deleted allele (*Smchd1*^−^) in trans to the *Smchd1* floxed (*Smchd1*^fl^) allele (49). The *Smchd1*^−^ allele was generated from the *Smchd1*^fl^ allele using a line of C57BL/6 mice expressing a constitutive Cre transgene (50). The *Smchd1*^fl^ and *Smchd1*^Del^ lines were produced and maintained on the C57BL/6 background. The Zp3-Cre line was backcrossed onto the *Smchd1*^fl^ allele (51).

The CAST/EiJ strain was purchased from the Jackson laboratories. To study loss of zygotic Smchd1 along with loss of oocyte Smchd1, we produced an F1 line of mice from CAST/EiJ strain dams mated with C57BL/6 *Smchd1*^Del/+^ sires. By mating with C57BL/6 *Smchd1*^Del/+^ dams or *Sm-chd1*^Del/fl^; MMTV-Cre^T/+^ dams we could generate embryos where on average half of the imprinted clusters would have a CAST/EiJ allele in trans to C57BL/6, (and therefore be informative for allele-specific analyses) and also be null for *Smchd1*.

The *Smchd1*^GFP^ allele was backcrossed to C57BL/6 for more than 10 generations, and kept as a homozygous breeding line as previously described(28).

Genotyping for the *Smchd1* alleles and sex chromosomes was done as previously described (28). The Cre transgenes were genotyped using a general Cre PCR as previously described (52).

All post-implantation embryos were generated via natural timed matings. All preimplantation embryos were generated following superovulation as previously described (53).

All illustrations of embryos were adapted from the atlas of embryonic development (54) https://creativecommons.org/licenses/by/3.0/.

### Embryo, placental and ovary dissections

The embryonic portion of the E14.5 placenta was dissected as previously described (29). The dissection was learnt using a GFP transgene that is transmitted from the sire, and therefore only present in the embryonic portion of the placenta. The yolk sac or a portion of the embryo was taken for genotyping, while the placental piece was snap frozen for later RNA and DNA preparation. The E9.5 embryos were dissected and snap frozen whole.

### Sectioning and immunofluorescence studies of ovary

The ovaries from 6-14-week-old mice were harvested and fixed in 10% formalin overnight. Immunofluorescence was performed on paraffin embedded, serially sectioned (4 μm) ovaries. A total of 5-9 slides (3 sections per slide) were assessed from each ovary. Briefly, paraffin sections were dewaxed in histolene and heat-induced epitope retrieval was performed in 10 mM sodium citrate buffer (pH 6). Sections were blocked in 10% donkey serum (Sigma Aldrich, D9663) in Tris-sodium chloride (TN) buffer with 3% Bovine Serum Albumin (BSA) (Sigma Aldrich, A9418). Sections were then incubated with buffer only (as negative controls) or primary antibodies for 24 hours at 4°C in TN with 1% BSA in the following dilutions: 1:100 SMCHD1 #5 or #8 (own derivation) and 1:500 CD117/c-kit Antibody (NOVUS af1356). Donkey antiGoat Alexa 488 (ThermoFisher Scientific, A-11055) and Donkey anti-Rat IgG DyLight 550 (Invitrogen, SA5-10027) secondary antibodies were applied at 1:500 in TN after washing. Slides were cover-slipped with ProLong™ Diamond Antifade Mountant with DAPI (Thermo Fisher Scientific, MA, USA, P36931) and imaged via a confocal Nikon Eclipse 90i (Nikon Corp., Tokyo, Japan) microscope. Images were processed and analysed using FIJI software (55). A minimum of 5 follicles for primordial, late secondary follicles and 2–3 antral follicles were tracked and imaged across multiple sections for each genotype.

### Native GFP imaging of preimplantation embryos

Preimplantation embryos were collected by flushing oviducts/uteri at 2-cell, 4-cell, 8-cell, 16-cell, 32-cell and blastocyst stages, at E1.5, E2.0, E2.25, E2.75, E3.5 as described previously (56). Embryos collected from *Smchd1*^GFP/GFP^ females crossed *Smchd1*^GFP/GFP^ males were used as positive controls and embryos from *Smchd1*^+/+^ parents were used as negative controls simultaneously with the test embryos to ensure autofluorescence could be accounted for in imaging for GFP. Preimplantation embryos were fixed with 4% paraformaldehyde for 10 min, washed with PBS and imaged in Fluorobrite DMEM media (ThermoFisher scientific A1896701) using an AxioObserver microscope (Zeiss) at 20x magnification. Images were processed and analysed using FIJI software (55). For each experiment at least 3 embryos were scored.

### Immunofluorescence of preimplantation embryos

Embryos were collected from *Smchd1*^GFP/GFP^ females crossed with *Smchd1*^+/+^ male and *Smchd1*^+/+^ females crossed with *Smchd1*^GFP/GFP^ male as tests, *Smchd1*^+/+^ crossed with *Smchd1*^+/+^ as negative controls, and *Smchd1*^GFP/GFP^ females crossed *Smchd1*^GFP/GFP^ male as positive controls at 8-cell, 12–16-cell and 24–32-cell stage embryos. The zona pellucida was removed using Acid tyrodes solution (Sigma T1788), then embryos washed with M2 solution (Sigma M7167) and fixed with 2% PFA for 20 min at room temperature. Embryos were permeabilized in 0.1% Triton-X100 (Sigma T8787) in PBS for 20 min at room temperature. Embryos were blocked in 0.25% gelatin in PBS (gelatin, Sigma, G1393) for 20 min at room temperature. Embryos were transferred to primary antibody, rabbit anti-GFP (Thermofisher A11122, lot 2015993) diluted in block and incubated for 1 hour. Embryos were washed with PBS then transferred to secondary antibody, anti-rabbit Alexa 488 (Thermofisher A21206, lot 1874771) diluted in block and incubated for 40 min in a dark humidified chamber. The embryos were finally washed with PBS, stained with DAPI for 10 min, washed with PBS again and mounted into Vectashield H-1000 (Vector labs). The embryos were imaged using the Zeiss LSM 880 system, 40x magnification with airyscan processing. Images were processed and analysed using FIJI software (55). Negative and positive controls were used to normalise fluorescence signal.

### Bulk RNA and DNA preparation from embryos and placentae

RNA and DNA were prepared from snap-frozen samples either using a Qiagen All prep kit (Qiagen, Cat # 80204), or a Zymo Quick DNA/RNA miniprep plus kit (Zymo research, Catalog # D7003). DNA and RNA were quantified using nanodrop (Denovix DS-11 spectrophotometer).

### Bulk RNA-sequencing and analysis

Libraries were prepared using TruSeq V1 or V2 RNA sample preparation kits from 500 ng total RNA as per manufacturers’ instructions. Fragments above 200 bp were size- selected and cleaned up using AMPure XP magnetic beads. Final cDNA libraries were quantified using D1000 tape on the TapeStation (4200, Agilent Technologies) for sequencing on the Illumina Nextseq500 platform using 80-bp, paired-end reads.

RNA-seq reads were trimmed for adapter and low quality sequences using TrimGalore! v0.4.4, before mapping onto the GRCm38 mouse genome reference N-masked for CAST/EiJ SNPs (prepared with SNPsplit v0.3.2 (57) with HISAT2 v2.0.5 (58), in paired-end mode and disabling soft-clipping. Alignments specific to the C57BL/6 and CAST/EiJ alleles were separated using SNPsplit v0.3.2 in paired-end mode.

Gene counts were obtained in R 3.5.1 (59) from the split and non-split bam files with the featureCounts function from the Rsubread package (1.32.1 (60, 61)), provided with the GRCm38.90 GTF annotation downloaded from Ensembl, and ignoring multi-mapping or multi-overlapping reads.

Because the samples for the maternal and zygotic deletion experiment are first-generation backcrosses of CastB6F1s to C57BL/6, differential expression between alleles can only be performed in genomic regions that are heterozygous (half of the genome, on average). To identify these regions for each sample, we fit a recursive partition tree with R rpart (59, 62, 63) function to the proportion of CAST/EiJ reads in each 100-kb bin tiling the genome, with options minsplit = 4, cp = 0.05.

Genes were defined as expressed and retained for differential expression analysis if they had at least one count per million (cpm) in at least a third of the libraries in the non-haplotyped data.

For analysis of global expression changes, total (non-haplotyped) counts were normalised in edgeR (3.24.0 (64, 65)) with the TMM method (66), and differential expression analysis performed with quasi-likelihood F-tests (67). P-values were corrected with the Benjamini-Hochberg method (68). Differential expression results were visualised with Glimma (1.10.0 (69)), with differential expression cut-offs of adjusted p-value < 0.05 and log2-fold-change > 1. Plots were generated in R with the ggplot2(70), cowplot (71), ggbeeswarm (72), ggrepel (73), ggrastr (74), and pheatmap (75) packages.

Testing for changes in imprinted expression was performed as follows: we compiled a list of all 316 known mouse imprinted genes and kept autosomal genes expressed in the experiment of interest (based on total counts, as above). To determine whether they were imprinted in the tissue of interest, we then fitted a logistic regression with the glm function from the stats package (59) on the maternal and paternal counts for each gene in the wild-type samples and retained the genes with an absolute log-ratio of expression greater than or equal to log(1.5). Genes without on average at least 10 haplotyped counts per sample for each genotype were also excluded from further analysis.

To investigate the effect of *Smchd1* oocyte-deletion on imprinted expression, we used edgeR’s paired sample design (76) with tagwise dispersion to model the maternal and paternal counts as a function of genotype (and Cre-deletion, in the case of placental samples where we had both Zp3-Cre and MMTV-Cre samples). We tested the effect of genotype with a likelihood ratio test. P-values were corrected with the Benjamini-Hochberg method. Genes were considered differentially imprinted when the adjusted p-value was less than 0.05 and the absolute difference in silent allele proportion average between wild-type and oocyte-deleted samples was greater than 5% (for at least one Cre construct in the case of placenta).

We could not use the same approach for the zygotic-and-oocyte deletion experiment as the genetic heterogeneity of the samples resulted in a haplotyped counts matrix with about half of the values missing. Instead, we fitted a beta-binomial regression with the betabin function from package the aod (77) package on the maternal and paternal counts for each gene with informative samples, with a fixed dispersion of 0.01 estimated from the oocyte-only deletion data. We used the same criteria of 5% false discovery (FDR) and 5% absolute difference in silent allele proportion to call differential imprinting.

### Bulk reduced representation bisulfite sequencing and analysis

DNA extracted above was cleaned up using Zymo research DNA Clean & concentrator-5 kit. DNA was quantified using Qubit dsDNA assay kit (ThermoFisher Scientific Q32853) and 100 ng were used as input for library preparation with the NuGEN Ovation RRBS methyl-seq system (Integrated sciences). Bisulfite conversion was carried out using QIAGEN EpiTect Fast DNA Bisulfite Kit. Quantitative and Qualitative analysis of library prep was carried out using D1000 tape on TapeStation 2200 (Agilent Technologies). Samples were sequenced on the HiSeq2500 platform using 100-bp paired-end reads.

Paired RRBS reads were trimmed first for adapter and low quality sequences with TrimGalore! v0.4.4 specifying the options -a AGATCGGAAGAGC -a2 AAATCAAAAAAAC, and then for the diversity bases introduced during library preparation with the trimRRBSdiversityAdaptCustomers.py script provided by NuGEN (https://github.com/nugentechnologies/NuMetRRBS). Trimmed reads were mapped onto the GRCm38 mouse genome reference N-masked for CAST/EiJ SNPs with Bismark v0.20.0 (78) and the alignments were split by allele with SNPsplit v0.3.2 in bisulfite mode. Methylation calls were extracted with Bismark’s `bismark_methylation_extractor` function.

We compiled a list of 45 known imprinted Differentially Methylated Regions (DMRs) and aggregated the haplotyped methylated and unmethylated counts over these regions. Regions with fewer than 10 counts per sample on average for wild-type and oocyte-deleted genotypes were filtered out, as were regions with an average methylation difference between the parental alleles in wild-type samples of less than 10%. Methylated and unmethylated counts for the hypermethylated allele were analysed with edgeR’s paired sample design, setting the dispersion as the common dispersion (76). P-values were adjusted for multiple testing with the Benjamini-Hochberg correction. Hypermethylated alleles were considered to be differentially methylated between wild-type and oocyte-deleted samples when the adjusted FDR was below 0.05.

### Smchd1-GFP ChIP-Seq in neural stem cells

Three primary female *Smchd1*^GFP/GFP^ NSC lines were derived as previously described (26) and harvested with Accutase (Sigma-Aldrich), washed with culture medium and PBS and cross-linked with 1% formaldehyde (vol/vol) for 10 min at room temperature with rotation, and subsequently quenched with glycine. The cells were immediately pelleted at 456 g for 5 minutes at 4°C and washed twice in cold PBS. The crosslinked pellets were then snap-frozen in dry ice.

For each cell line, 4 × 10^7^ crosslinked nuclei were extracted from frozen pellets by incubating on ice for 10 minutes in 14 mL of ChIP buffer (150 mM NaCl, 50 mM Tris-HCl pH 7.5, 5 mM EDTA, 0.5% vol/vol Igepal CA-630, 1% Triton X-100, 1X cOmplete cocktail (Roche) and homogenizing 25 times in a tight dounce on ice. Nuclei were pelleted at 12,000 g for 1 min at 4°C, washed with ChIP buffer then resuspended in 1.6 mL of of MNase buffer with 1x BSA (NEB). The nuclei solution was preincubated at 37°C for 5 minutes, then 2 × 10^4^ U of MNase (NEB) was added and incubated for a further 5 minutes. The reaction was stopped by adding 10 mM of EDTA and incubating on ice for 10 minutes. Nuclei were pelleted (4°C, 12,000 g, 1 minute), resuspended in 520 μL of ChIP buffer then sonicated with a Covaris S220 sonicator (peak power, 125; duty factor, 10; cycle/burst, 200; duration, 15 s) in Covaris microTubes. Sonicated solution was diluted 10 times with ChIP buffer then spun at 12,000 g at 4°C for 1 minute to clear debris. 20 ul of the supernatant was taken for the whole cell extract (WCE) samples and the rest was used for immunoprecipitation overnight at 4°C with rotation with 16 μg of anti-GFP antibody (Invitrogen A11122). The chromatin was then cleared by centrifugation (12,000 g for 10 minutes at 4°C) and 80 μL of protein G DynaBeads (ThermoFisher, washed 3 times in cold ChIP buffer right before use) were added before incubating at 4°C for 1 hour with rotation. The samples were then washed 6 times with cold ChIP buffer. The antibody-bound chromatin was eluted from the beads with two rounds of 400 μL of Elution buffer (1% SDS, 0.1 M NaHCO3) by rotating at room temperature for 15 minutes. 8 μL of 5 M NaCL and 1 μL of RNase A (NEB) were added to every 200 μL of eluate and to the WCE samples (diluted to 200 μL with Elution buffer), incubated overnight at 65°C to reverse crosslinking then treated with 4 μL of 20 μg/mL Proteinase K at 65°C for 1 hour. DNA was extracted with Zymo ChIP DNA clean and concentrator kit. Libraries were generated with an Illumina TruSeq DNA Sample Preparation Kit. 200- to 400-bp fragments were size-selected with AMPure XP magnetic beads. Libraries were quantified with a D1000 tape in a 4200 Tapestation (Agilent). Libraries were pooled and sequenced on the Illumina NextSeq platform, with 75-bp single-end reads.

Adapter trimming was performed with Trim galore! v0.4.4, library QC was assessed with FastQC v0.11.8(79) before mapping with Bowtie2 v2.3.4.1(80) with default options to the GRCm38.p6 version of the reference mouse genome. BAM files were imported into SeqMonk v1.45.1(81) extending them by 150 bp and peaks were called with the MACS-style caller within the SeqMonk package (settings for 300-bp fragments, *p* < 1 × 10^*−*5^) by merging all three Smchd1GFP and both WCE biological replicates into replicate sets. ChIP-seq tracks were produced with SeqMonk by defining probes with a running window (1000 bp width; 250 bp step), doing a read-count quantitation then normalizing by library size before doing a match distribution normalization within each replicate set and smoothing over 5 adjacent probes.

### Single-preimplantation embryo M&T-seq

E2.75 embryos were sorted into a preprepared lysis buffer containing 2.5 μL of RLT Plus (Qiagen) with 1 U/μL SUPERase-IN (Ambion). Genomic DNA and mRNA were separated using oligo-dT conjugated magnetic beads according to the G&T-seq protocol (82), however gDNA was eluted into 10 μL of H2O and 16 cycles of amplification followed for cDNA synthesis.

Bisulfite libraries were prepared using an adapted scBS-seq protocol (83). Briefly, bisulfite conversion was performed before introducing Illumina adaptor sequences with random priming oligos. Only one round of first and second strand synthesis was performed, using primers compatible with the NEBNext multiplex oligos for library production (Forward: 5’-CTACACGACGCTCTTCCGATCTNNNNNN-3’; Reverse: 5’-CAGACGTGTGCTCTTCCGATCTNNNNNN-3’). Sixteen cycles of library amplification followed and all Ampure XP bead (Beckman Coulter) purifications were performed at 0.65×. To assess library quality and quantity a High-Sensitivity D5000 ScreenTape on the Agilent TapeStation and the ProNex NGS Library Quantification Kit were used. Single-embryo bisulfite libraries were pooled for 150-bp paired-end sequencing on a NovaSeq6000.

RNA-seq libraries were prepared from amplified cDNA using the Nextera XT kit (Illumina) according to the manufacturer’s guidelines but using one-fifth volumes. Single-embryo RNA-seq libraries were pooled for 75-bp paired-end sequencing on a NextSeq500.

### Single preimplantation embryo M&T-seq analysis

#### Methylome analysis

To remove sequence biases due to the post-bisulfite adapter tagging, the first nine base pairs from read 1 and read 2 were removed with Trim Galore! v0.4.4 (http://www.bioinformatics.babraham.ac.uk/projects/trim_galore/), in addition to adapter and quality trimming, with options (`--clip_R1 9 --clip_R2 9`). The reads were then first mapped to the CAST/EiJ-masked GRCm38 genome in paired-end mode with Bismark v0.20.0 with the `--pbat` option to allow dovetail mapping, and keeping unmapped reads (`--unmapped`). Unmapped reads 1 and 2 were then mapped in singleend mode, with the `--pbat` option set for read 1. Each alignment file (paired, unmapped reads 1 and unmapped reads 2) was then deduplicated with `deduplicate_bismark`, and haplotyped with SNPsplit v0.3.2. Methylation calls were obtained with `bismark_methylation_extractor` and pooled (paired, unmapped reads 1 and unmapped reads 2).

Embryos were sexed based on their CAST/EiJ-haplotyped bisulfite reads mapping to the X chromosome. Embryos that did not produce a library, showed evidence of maternal contamination (much higher proportion of C57BL/6 reads compared to CAST/EiJ reads in the methylome and transcriptome data), or displayed chromosomal abnormalities were excluded.

Differences in methylation at known imprinted DMRs were tested in edgeR as for the bulk samples, although we also tested the non-split data in additition to the hypermethylated allele to increase the coverage and have more testable regions.

#### Transcriptome analysis

RNA-seq analyses for the single embryos passing quality control were performed as for the bulk samples, except that the dispersion was set as the common dispersion (rather than tagwise) to avoid unreliable estimates of dispersion for individual genes on sparser single-embryo expression data.

## DATA AVAILABILITY

All the sequencing data for this study are available from the Short Read Archive under the BioProject accession PRJNA530651. List of imprinted genes, imprinted DMRs and results of the genome-wide expression analyses can be consulted on http://bioinf.wehi.edu.au/smchd1_mat_effects/.

## ETHICS APPROVAL AND CONSENT

All mice were housed and bred at The Walter and Eliza Hall Institute of Medical Research, under ethics approval from the The Walter and Eliza Hall Institute of Medical Research Animal Ethics Committee (approval numbers 2014.026, 2018.004).

## AUTHOR CONTRIBUTIONS

IW, QG and MEB designed the experiments, analysed and interpreted the data and wrote the manuscript. IW performed all microscopy experiments and most embryonic dissections. QG analysed all RNAseq, allele-specific RNAseq, allele-specific RRBS and M&Tseq data. SAK performed dissections and analysed some genomic data and contributed to interpretation of data and editing the manuscript. ATdF performed ChIP-seq and its analysis and edited the manuscript. JS performed the ovary section immunofluorescence and analysed this data. AK analysed some genomics experiments and contributed to interpretation of data. TB and KB performed dissections and made genomics libraires. KH contributed to experimental design and interpretation of the ovary section experiments. HJL contributed to experimental design of the single preimplantation embryo M&Tseq. MER contributed to design and analysis of all genomics experiments. SAK, AtdF, AK and MER edited the manuscript. All authors read and approved the final manuscript.

## ACKNOWLEDGEMENTS

We thank Kay-Uwe Wagner, Jane Visvader, Patrick Western and Graham Kay for provision of the MMTV-Cre, Zp3-Cre and Smchd1-GFP mice used in the study. We thank Stephen Wilcox and Jafar Jabbari for sequencing. We thank Peter Hickey, Yunshun Chen and Gordon Smyth for statistical advice and Natasha Jansz for critical reading of the manuscript.

## Supplementary figures

**Fig. S1.**
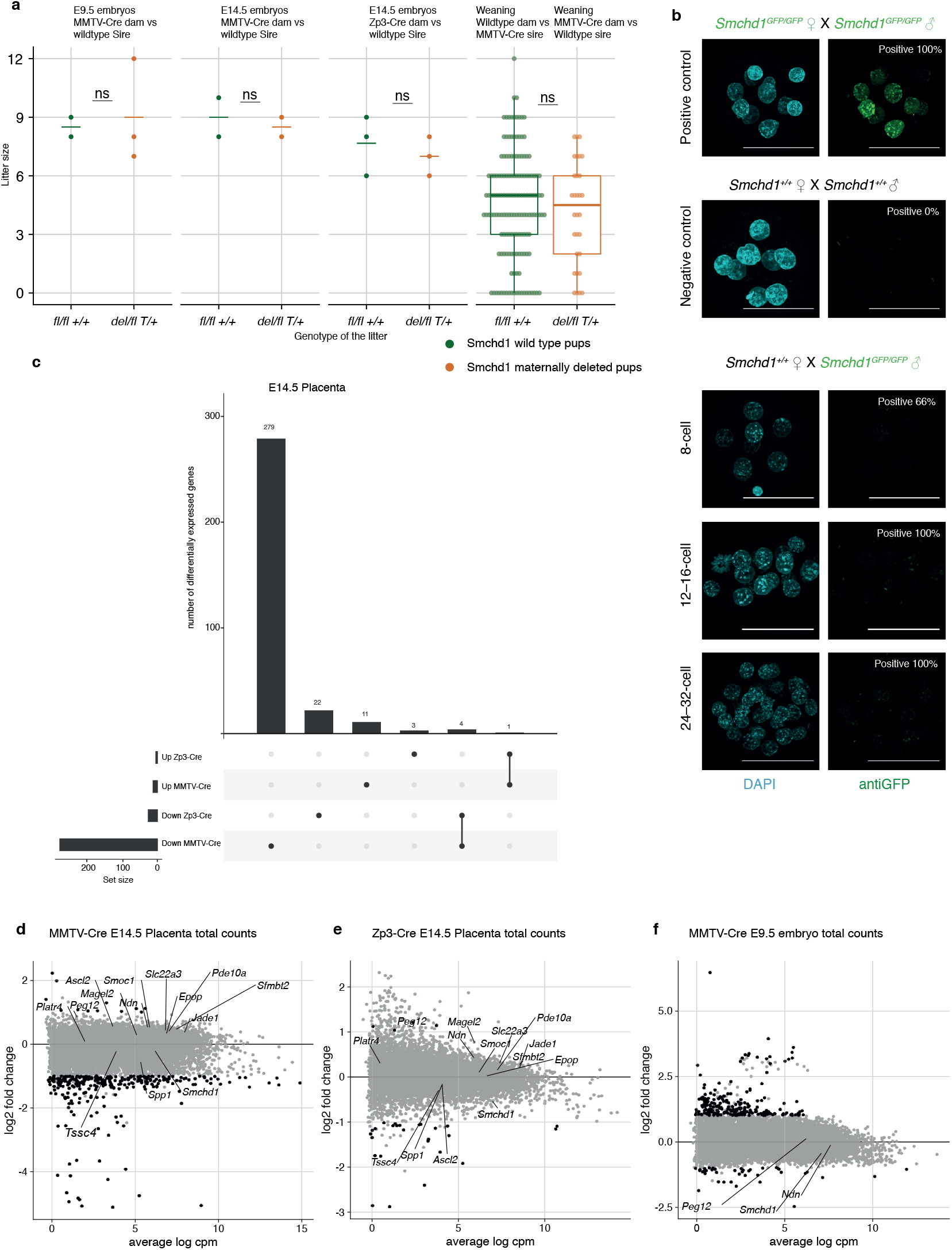
**a.** Embryo lethality data for Smchd1 maternal null embryos and animals at weaning, compared to control litters generated from *Smchd1*^fl/fl^ non-transgenic dams, or a reciprocal cross. The p-value was calculated using student two-tailed T-test. **b.** Immunofluorescence detection of maternal Smchd1-GFP in the nuclei of 8-cell embryos, with *Smchd1*^+/+^ used as negative controls and *Smchd1*^GFP/GFP^ embryos were used as positive controls. Zygotic Smchd1 was detectable in some cells from the 8-cell stage onwards but comparatively less signal intensity to the GFP control when embryos were imaged with identical settings, in experiments performed on the same day. n = 2 embryos. **c.** Intersection of the differentially expressed gene lists in MMTV-Cre and Zp3-Cre maternal deletion experiments, E14.5 placental samples. *d-f*. MA-plots of total gene expression in MMTV-Cre (**e**) and Zp3-Cre (**d**) maternal deletion experiments, E14.5 placental samples, and MMTV-Cre E9.5 embryos (**f**). Genes below the 5% FDR and differentially expressed by at least 2-fold are plotted in black. *Smchd1* and genes with partial loss of imprinting are labelled.

**Fig. S2.**
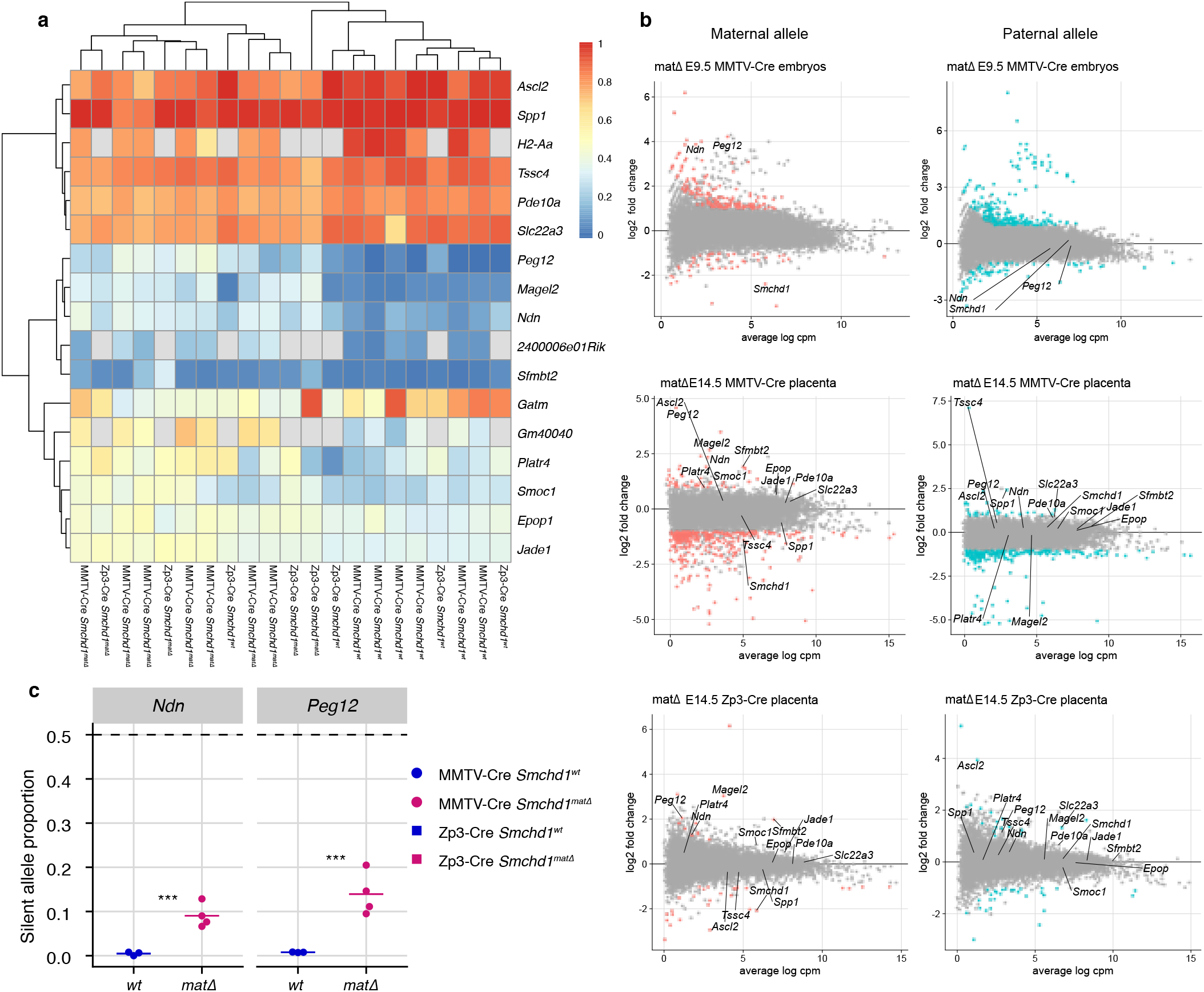
**a.** Heatmap of maternal allele expression proportion for genes that are differentially imprinted in at least one experimental set (MMTV-Cre and Zp3-Cre E14.5 placenta) **b.** MA-plots of allelic gene expression in MMTV-Cre E9.5 embryos and E14.5 placentae, and Zp3-Cre E14.5 placentae. Genes below the 5% FDR and differentially expressed by at least 2-fold are plotted in colour. Smchd1 and genes with partial loss of imprinting are labelled. **c.** Expression of the silent allele as a proportion of total expression of the gene at the *Snprn* cluster genes *Ndn* and *Peg12*, in E9.5 *Smchd1* maternal null embryos. * *p* < 0.05, ** *p* < 0.01, *** *p* < 0.001, when the difference in silent allele proportions is at least 5%.

**Fig. S3.**
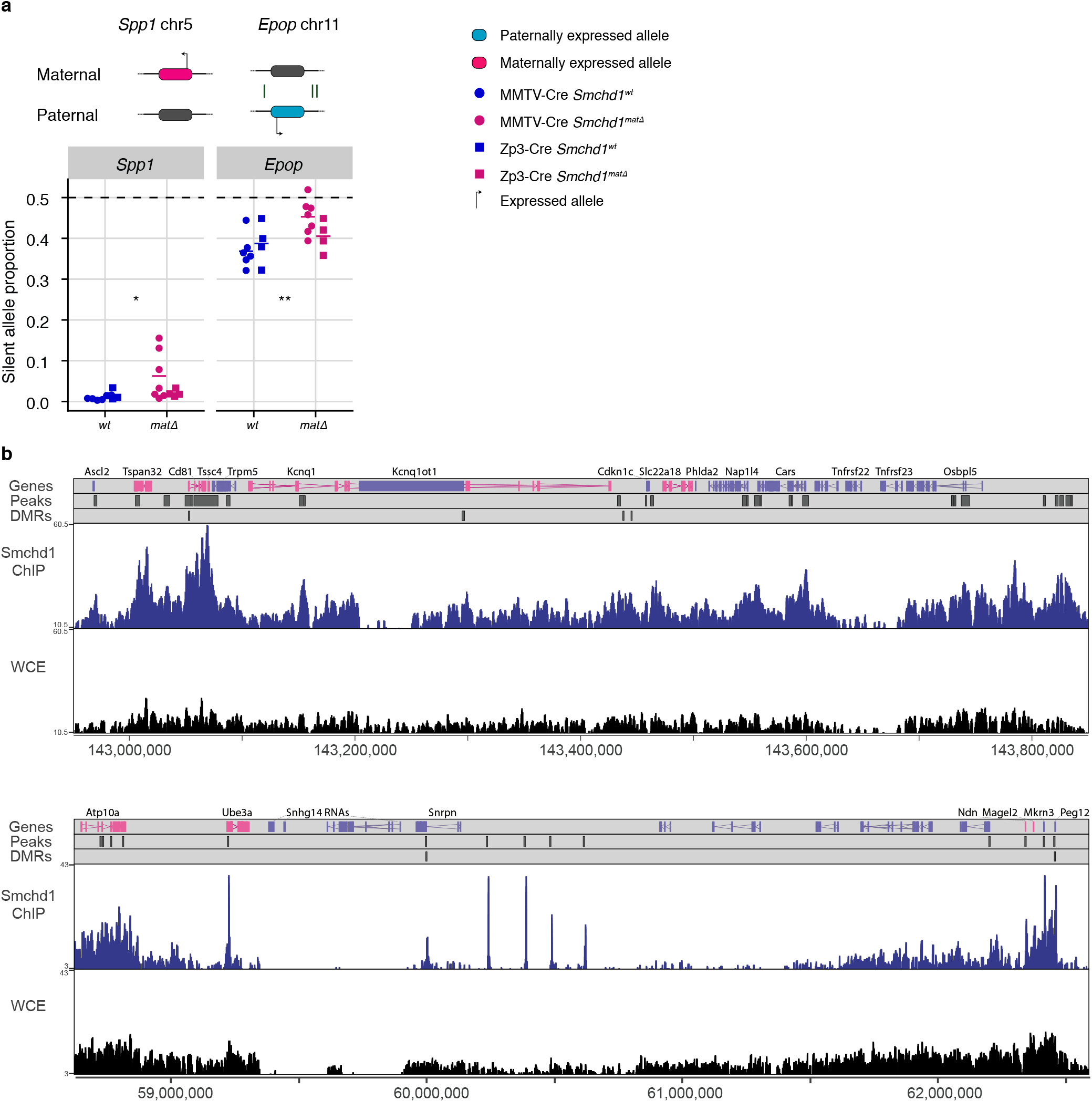
**a.** Expression of the silent allele as a proportion of total expression of the gene at the lone imprinted genes *Spp1* and *Epop*, in the embryonic portion of the placenta of *Smchd1* maternal null conceptuses at E14.5. ns *p >* 0.05, * *p* < 0.05, ** *p* < 0.01, *** *p* < 0.001, when the difference in silent allele proportions is at least 5%. **b.**ChIP-seq for Smchd1-GFP over the *Kcnq1* (upper panel, chr7:142950000–143850000) and *Snrpn* (lower panel chr7:58625000–62593000) gene clusters. Gene tracks showing mRNA annotation are on top (some annotated genes not mentioned in this study were removed for clarity). MACS2-like Smchd1-GFP ChIP-seq peaks are shown as dark grey rectangles, as are differentially methylated regions in the track below. Smoothed Smchd1-GFP (average of 3 biological replicates) and whole cell extract (WCE) (average of 2 biological replicates) tracks show normalised enrichment of reads from *Smchd1*^GFP/GFP^ neural stem cells. Scale is normalised ChIP-seq reads and the y-axis starts at a value equal to 75% of the mean value of the WCE of the region shown.

**Fig. S4.**
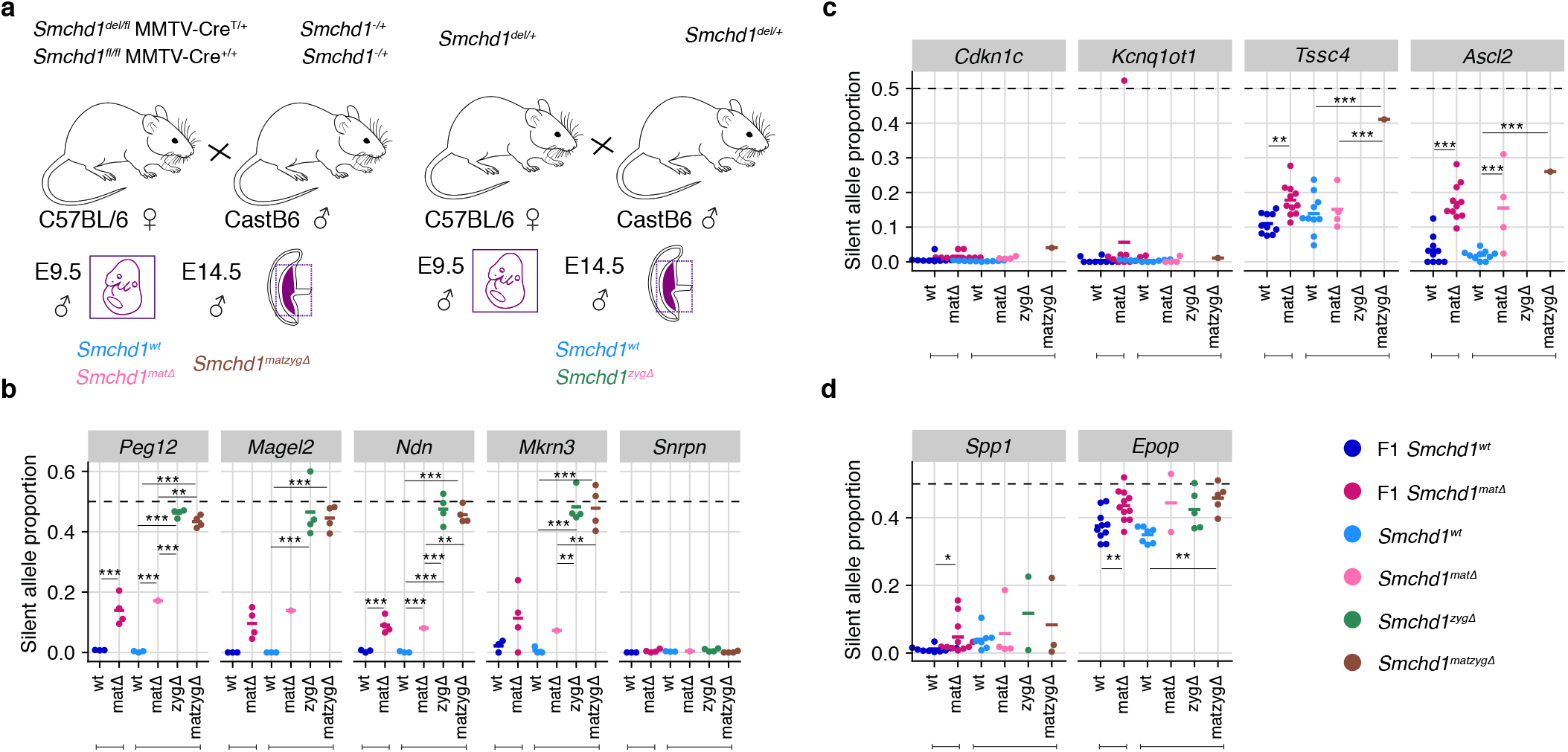
**a.** Schematic for breeding scheme to generate oocyte (matΔ), zygotic (zygΔ), oocyte-and-zygotic deletion (matzygΔ) of Smchd1. Strain background is shown underneath each parent. Genotype is shown above each parent. Allele-specific RNA-seq expression profiles of the silent allele is shown as a proportion of total expression at the **b.** Snrpn cluster in E9.5 embryo **c.** *Kcnq1* cluster and **d.** *Spp1* and *Epop* in the embryonic portion of the placenta at E14.5. * *p* < 0.05, ** *p* < 0.01, *** *p* < 0.001, when the difference in silent allele proportions is at least 5%.

**Fig. S5.**
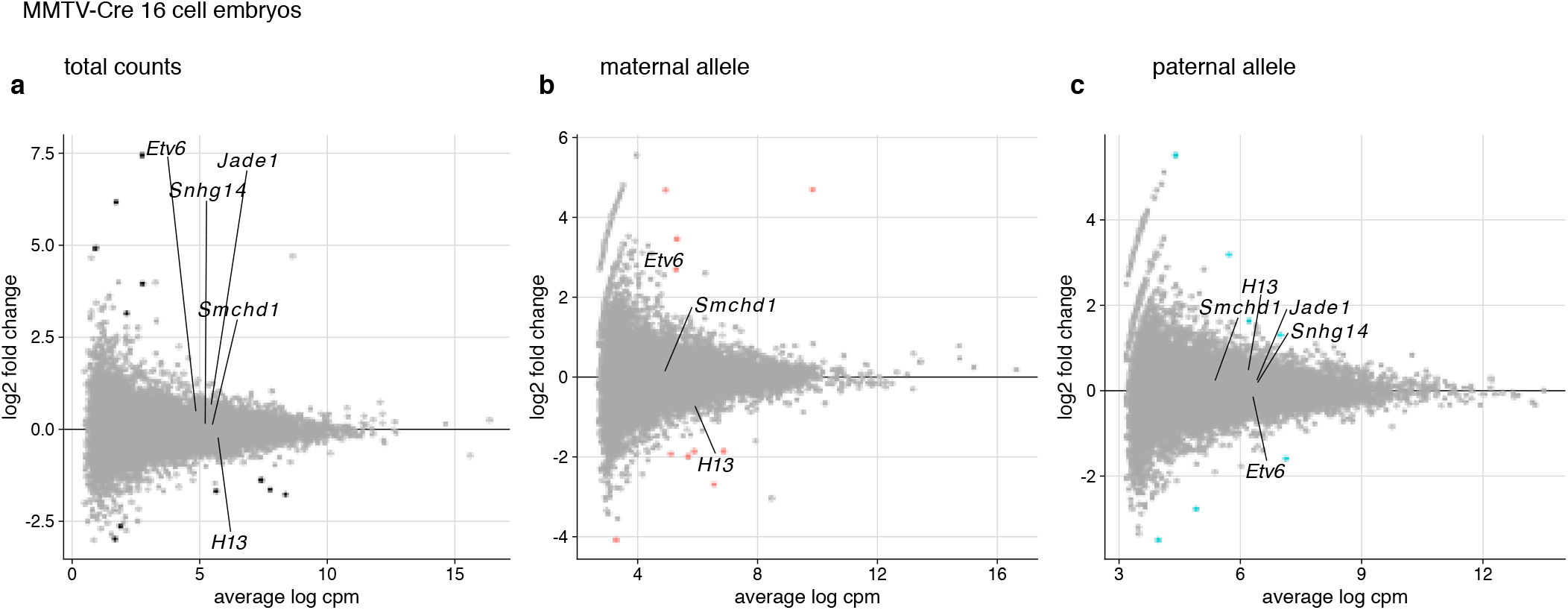
MA-plots of total (**a**) and allelic (**b**,**c**) gene expression in MMTV-Cre maternal deletion E2.75 embryos experiments. Genes below the 5% FDR and differentially expressed by at least 2-fold are plotted in black. *Smchd1* and genes with partial loss of imprinting are labelled.

## Bibliography

1. James McGrath and Davor Solter. Completion of mouse embryogenesis requires both the maternal and paternal genomes. Cell, 37(1):179–183, 1984. ISSN 00928674. doi: 10.1016/0092-8674(84)90313-1.

2. M. A. H. Surani, S. C. Barton, and M. L. Norris. Development of reconstituted mouse eggs suggests imprinting of the genome during gametogenesis. Nature, 308(5959): 548–550, 1984. ISSN 0028-0836 1476-4687. doi: 10.1038/308548a0.

3. T. Kishino, M. Lalande, and J. Wagstaff. UBE3A/E6-AP mutations cause Angelman syndrome. Nature Genetics, 15(1):70–3, 1997. ISSN 1061-4036 (Print) 1061-4036 (Linking). doi: 10.1038/ng0197-70.

4. T. Matsuura, J. S. Sutcliffe, P. Fang, R. J. Galjaard, Y. H. Jiang, C. S. Benton, J. M. Rommens, and A. L. Beaudet. De novo truncating mutations in E6-AP ubiquitin-protein ligase gene (UBE3A) in Angelman syndrome. Nature Genetics, 15(1):74–7, 1997. ISSN 1061-4036 (Print) 1061-4036 (Linking). doi: 10.1038/ng0197-74.

5. R. D. Nicholls, J. H. Knoll, M. G. Butler, S. Karam, and M. Lalande. Genetic imprinting suggested by maternal heterodisomy in nondeletion Prader-Willi syndrome. Nature, 342 (6247):281–5, 1989. ISSN 0028-0836 (Print) 0028-0836 (Linking). doi: 10.1038/342281a0.

6. Jo Peters. The role of genomic imprinting in biology and disease: an expanding view. Nature Reviews Genetics, 15(8):517–530, 2014. ISSN 1471-0064. doi: 10.1038/nrg3766.

7. F. L. Sun, W. L. Dean, G. Kelsey, N. D. Allen, and W. Reik. Transactivation of Igf2 in a mouse model of Beckwith-Wiedemann syndrome. Nature, 389(6653):809–15, 1997. ISSN 0028-0836 (Print) 0028-0836 (Linking). doi: 10.1038/39797.

8. J. S. Sutcliffe, Y. H. Jiang, R. J. Galijaard, T. Matsuura, P. Fang, T. Kubota, S. L. Christian, J. Bressler, B. Cattanach, D. H. Ledbetter, and A. L. Beaudet. The E6-Ap ubiquitin-protein ligase (UBE3A) gene is localized within a narrowed Angelman syndrome critical region. Genome Research, 7(4):368–77, 1997. ISSN 1088-9051 (Print) 1088-9051 (Linking). doi: 10.1101/gr.7.4.368.

9. V. Tucci, A. R. Isles, G. Kelsey, A. C. Ferguson-Smith, and Group Erice Imprinting. Genomic imprinting and physiological processes in mammals. Cell, 176(5):952–965, 2019. ISSN 1097-4172 (Electronic) 0092-8674 (Linking). doi: 10.1016/j.cell.2019.01.043.

10. P. Zhang, N. J. Liegeois, C. Wong, M. Finegold, H. Hou, J. C. Thompson, A. Silverman, J. W. Harper, R. A. DePinho, and S. J. Elledge. Altered cell differentiation and proliferation in mice lacking p57KIP2 indicates a role in Beckwith-Wiedemann syndrome. Nature, 387(6629):151–8, 1997. ISSN 0028-0836 (Print) 0028-0836 (Linking). doi: 10.1038/387151a0.

11. D. P. Barlow and M. S. Bartolomei. Genomic imprinting in mammals. Cold Spring Harbour Perspectives in Biology, 6(2), 2014. ISSN 1943-0264 (Electronic) 1943-0264 (Linking). doi: 10.1101/cshperspect.a018382.

12. A. C. Ferguson-Smith. Genomic imprinting: the emergence of an epigenetic paradigm. Nature Reviews Genetics, 12(8):565–75, 2011. ISSN 1471-0064 (Electronic) 1471-0056 (Linking). doi: 10.1038/nrg3032.

13. A. Inoue, Z. Chen, Q. Yin, and Y. Zhang. Maternal Eed knockout causes loss of H3K27me3 imprinting and random X inactivation in the extraembryonic cells. Genes & Development, 32(23-24):1525–1536, 2018. ISSN 1549-5477 (Electronic) 0890-9369 (Linking). doi: 10.1101/gad.318675.118.

14. Azusa Inoue, Lan Jiang, Falong Lu, and Yi Zhang. Genomic imprinting of xist by maternal h3k27me3. Genes & Development, 31(19):1927–1932, 2017. ISSN 0890-9369 1549-5477. doi: 10.1101/gad.304113.117.

15. K. A. Alexander, X. Wang, M. Shibata, A. G. Clark, and M. J. Garcia-Garcia. TRIM28 controls genomic imprinting through distinct mechanisms during and after early genome-wide reprogramming. Cell Reports, 13(6):1194–1205, 2015. ISSN 2211-1247 (Electronic). doi: 10.1016/j.celrep.2015.09.078.

16. H. Hiura, Y. Obata, J. Komiyama, M. Shirai, and T. Kono. Oocyte growth-dependent progression of maternal imprinting in mice. Genes Cells, 11(4):353–61, 2006. ISSN 1356-9597 (Print) 1356-9597 (Linking). doi: 10.1111/j.1365-2443.2006.00943.x.

17. Carina Y. Howell, Timothy H. Bestor, Feng Ding, Keith E. Latham, Carmen Mertineit, Jacquetta M. Trasler, and J. Richard Chaillet. Genomic imprinting disrupted by a maternal effect mutation in the Dnmt1 gene. Cell, 104(6):829–838, 2001. ISSN 00928674. doi: 10.1016/s0092-8674(01)00280-x.

18. X. Li, M. Ito, F. Zhou, N. Youngson, X. Zuo, P. Leder, and A. C. Ferguson-Smith. A maternal-zygotic effect gene, zfp57, maintains both maternal and paternal imprints. Developmental Cell, 15(4):547–57, 2008. ISSN 1878-1551 (Electronic) 1534-5807 (Linking). doi: 10.1016/j.devcel.2008.08.014.

19. S. Mahadevan, V. Sathappan, B. Utama, I. Lorenzo, K. Kaskar, and I. B. Van den Veyver. Maternally expressed NLRP2 links the subcortical maternal complex (SCMC) to fertility, embryogenesis and epigenetic reprogramming. Scientific Reports, 7:44667, 2017. ISSN 2045-2322 (Electronic) 2045-2322 (Linking). doi: 10.1038/srep44667.

20. S. McGraw, C. C. Oakes, J. Martel, M. C. Cirio, P. de Zeeuw, W. Mak, C. Plass, M. S. Bartolomei, J. R. Chaillet, and J. M. Trasler. Loss of DNMT1o disrupts imprinted X chromosome inactivation and accentuates placental defects in females. PLoS Genetics, 9(11):e1003873, 2013. ISSN 1553-7404 (Electronic) 1553-7390 (Linking). doi: 10.1371/journal.pgen.1003873.

21. D. M. Messerschmidt, W. de Vries, M. Ito, D. Solter, A. Ferguson-Smith, and B. B. Knowles. Trim28 is required for epigenetic stability during mouse oocyte to embryo transition. Science, 335(6075):1499–502, 2012. ISSN 1095-9203 (Electronic) 0036-8075 (Linking). doi: 10.1126/science.1216154.

22. Bernhard Payer, Mitinori Saitou, Sheila C. Barton, Rosemary Thresher, John P. C. Dixon, Dirk Zahn, William H. Colledge, Mark B. L. Carlton, Toru Nakano, and M. Azim Surani. Stella is a maternal effect gene required for normal early development in mice. Current Biology, 13(23):2110–2117, 2003. ISSN 09609822. doi: 10.1016/j.cub.2003.11.026.

23. J. Shin, M. Bossenz, Y. Chung, H. Ma, M. Byron, N. Taniguchi-Ishigaki, X. Zhu, B. Jiao, L. L. Hall, M. R. Green, S. N. Jones, I. Hermans-Borgmeyer, J. B. Lawrence, and I. Bach. Maternal Rnf12/RLIM is required for imprinted X-chromosome inactivation in mice. Nature, 467(7318):977–81, 2010. ISSN 1476-4687 (Electronic) 0028-0836 (Linking). doi: 10.1038/nature09457.

24. M. E. Blewitt, A. V. Gendrel, Z. Y. Pang, D. B. Sparrow, N. Whitelaw, J. M. Craig, A. Apedaile, D. J. Hilton, S. L. Dunwoodie, N. Brockdorff, G. F. Kay, and E. Whitelaw. SmcHD1, containing a structural-maintenance-of-chromosomes hinge domain, has a critical role in X inactivation. Nature Genetics, 40(5):663–669, 2008. ISSN 1061-4036. doi: 10.1038/ng.142.

25. A. V. Gendrel, A. Apedaile, H. Coker, A. Termanis, I. Zvetkova, J. Godwin, Y. A. Tang, D. Huntley, G. Montana, S. Taylor, E. Giannoulatou, E. Heard, I. Stancheva, and N. Brockdorff. Smchd1-dependent and -independent pathways determine developmental dynamics of CpG island methylation on the inactive X chromosome. Developmental Cell, 23(2):265–79, 2012. ISSN 1878-1551 (Electronic) 1534-5807 (Linking). doi: 10.1016/j.devcel.2012.06.011.

26. K. Chen, J. Hu, D. L. Moore, R. Liu, S. A. Kessans, K. Breslin, I. S. Lucet, A. Keniry, H. S. Leong, C. L. Parish, D. J. Hilton, R. J. Lemmers, S. M. van der Maarel, P. E. Czabotar, R. C. Dobson, M. E. Ritchie, G. F. Kay, J. M. Murphy, and M. E. Blewitt. Genome-wide binding and mechanistic analyses of Smchd1-mediated epigenetic regulation. PNAS, 112(27):E3535–44, 2015. ISSN 1091-6490 (Electronic) 0027-8424 (Linking). doi: 10.1073/pnas.1504232112.

27. A. V. Gendrel, Y. A. Tang, M. Suzuki, J. Godwin, T. B. Nesterova, J. M. Greally, E. Heard, and N. Brockdorff. Epigenetic functions of Smchd1 repress gene clusters on the inactive X chromosome and on autosomes. Molecular Cell Biology, 33(16): 3150–65, 2013. ISSN 1098-5549 (Electronic) 0270-7306 (Linking). doi: 10.1128/MCB.00145-13.

28. Natasha Jansz, Andrew Keniry, Marie Trussart, Heidi Bildsoe, Tamara Beck, Ian D. Tonks, Arne W. Mould, Peter Hickey, Kelsey Breslin, Megan Iminitoff, Matthew E. Ritchie, Edwina McGlinn, Graham F. Kay, James M. Murphy, and Marnie E. Blewitt. Smchd1 regulates long-range chromatin interactions on the inactive X chromosome and at Hox clusters. Nature Structural & Molecular Biology, 2018. ISSN 1545-9993 1545-9985. doi: 10.1038/s41594-018-0111-z.

29. A. W. Mould, Z. Y. Pang, M. Pakusch, I. D. Tonks, M. Stark, D. Carrie, P. Mukhopadhyay, A. Seidel, J. J. Ellis, J. Deakin, M. J. Wakefield, L. Krause, M. E. Blewitt, and G. F. Kay. Smchd1 regulates a subset of autosomal genes subject to monoallelic expression in addition to being critical for X inactivation. Epigenetics & Chromatin, 6, 2013. ISSN 1756-8935. doi: Artn1910.1186/1756-8935-6-19.

30. N. Jansz, K. Chen, J. M. Murphy, and M. E. Blewitt. The epigenetic regulator SM-CHD1 in development and disease. Trends in Genetics, 33(4):233–243, 2017. ISSN 0168-9525 (Print) 0168-9525 (Linking). doi: 10.1016/j.tig.2017.01.007.

31. R. J. Lemmers, R. Tawil, L. M. Petek, J. Balog, G. J. Block, G. W. Santen, A. M. Amell, P. J. van der Vliet, R. Almomani, K. R. Straasheijm, Y. D. Krom, R. Klooster, Y. Sun, J. T. den Dunnen, Q. Helmer, C. M. Donlin-Smith, G. W. Padberg, B. G. van Engelen, J. C. de Greef, A. M. Aartsma-Rus, R. R. Frants, M. de Visser, C. Desnuelle, S. Sacconi, G. N. Filippova, B. Bakker, M. J. Bamshad, S. J. Tapscott, D. G. Miller, and S. M. van der Maarel. Digenic inheritance of an SMCHD1 mutation and an FSHD-permissive D4Z4 allele causes facioscapulohumeral muscular dystrophy type 2. Nature Genetics, 44(12):1370–4, 2012. ISSN 1546-1718 (Electronic) 1061-4036 (Linking). doi: 10.1038/ng.2454.

32. C. T. Gordon, S. Xue, G. Yigit, H. Filali, K. Chen, N. Rosin, K. I. Yoshiura, M. Oufa-dem, T. J. Beck, R. McGowan, A. C. Magee, J. Altmuller, C. Dion, H. Thiele, A. D. Gurzau, P. Nurnberg, D. Meschede, W. Muhlbauer, N. Okamoto, V. Varghese, R. Irving, S. Sigaudy, D. Williams, S. F. Ahmed, C. Bonnard, M. K. Kong, I. Ratbi, N. Fejjal, M. Fikri, S. C. Elalaoui, H. Reigstad, C. Bole-Feysot, P. Nitschke, N. Ragge, N. Levy, G. Tuncbilek, A. S. Teo, M. L. Cunningham, A. Sefiani, H. Kayserili, J. M. Murphy, C. Chatdokmaiprai, A. M. Hillmer, D. Wattanasirichaigoon, S. Lyonnet, F. Magdinier, A. Javed, M. E. Blewitt, J. Amiel, B. Wollnik, and B. Reversade. De novo mutations in SMCHD1 cause Bosma arhinia microphthalmia syndrome and abrogate nasal development. Nature Genetics, 49(2):249–255, 2017. ISSN 1546-1718 (Electronic) 1061-4036 (Linking). doi: 10.1038/ng.3765.

33. A. D. Gurzau, K. Chen, S. Xue, W. Dai, I. S. Lucet, T. T. N. Ly, B. Reversade, M. E. Blewitt, and J. M. Murphy. FSHD2- and BAMS-associated mutations confer opposing effects on SMCHD1 function. Journal of Biological Chemistry, 2018. ISSN 1083-351X (Electronic) 0021-9258 (Linking). doi: 10.1074/jbc.RA118.003104.

34. N. D. Shaw, H. Brand, Z. A. Kupchinsky, H. Bengani, L. Plummer, T. I. Jones, S. Erdin, K. A. Williamson, J. Rainger, A. Stortchevoi, K. Samocha, B. B. Currall, D. S. Dunican, R. L. Collins, J. R. Willer, A. Lek, M. Lek, M. Nassan, S. Pereira, T. Kammin, D. Lucente, A. Silva, C. M. Seabra, C. Chiang, Y. An, M. Ansari, J. K. Rainger, S. Joss, J. C. Smith, M. F. Lippincott, S. S. Singh, N. Patel, J. W. Jing, J. R. Law, N. Ferraro, A. Verloes, A. Rauch, K. Steindl, M. Zweier, I. Scheer, D. Sato, N. Okamoto, C. Jacobsen, J. Tryggestad, S. Chernausek, L. A. Schimmenti, B. Brasseur, C. Cesaretti, J. E. Garcia-Ortiz, T. P. Buitrago, O. P. Silva, J. D. Hoffman, W. Muhlbauer, K. W. Ruprecht, B. L. Loeys, M. Shino, A. M. Kaindl, C. H. Cho, C. C. Morton, R. R. Meehan, V. van Heyningen, E. C. Liao, R. Balasubramanian, J. E. Hall, S. B. Seminara, D. Macarthur, S. A. Moore, K. I. Yoshiura, J. F. Gusella, J. A. Marsh, Jr. Graham, J. M., A. E. Lin, N. Katsanis, P. L. Jones, Jr. Crowley, W. F., E. E. Davis, D. R. FitzPatrick, and M. E. Talkowski. SMCHD1 mutations associated with a rare muscular dystrophy can also cause isolated arhinia and Bosma arhinia microphthalmia syndrome. Nature Genetics, 49(2):238–248, 2017. ISSN 1546-1718 (Electronic) 1061-4036 (Linking). doi: 10.1038/ng.3743.

35. M. R. Gdula, T. B. Nesterova, G. Pintacuda, J. Godwin, Y. Zhan, H. Ozadam, M. McClellan, D. Moralli, F. Krueger, C. M. Green, W. Reik, S. Kriaucionis, E. Heard, J. Dekker, and N. Brockdorff. The non-canonical SMC protein SmcHD1 antagonises TAD formation and compartmentalisation on the inactive X chromosome. Nature Communications, 10(1):30, 2019. ISSN 2041-1723 (Electronic) 2041-1723 (Linking). doi: 10.1038/s41467-018-07907-2.

36. C. Y. Wang, T. Jegu, H. P. Chu, H. J. Oh, and J. T. Lee. SMCHD1 merges chromosome compartments and assists formation of super-structures on the inactive X. Cell, 174(2):406–421 e25, 2018. ISSN 1097-4172 (Electronic) 0092-8674 (Linking). doi: 10.1016/j.cell.2018.05.007.

37. U. Midic, K. A. Vincent, K. Wang, A. Lokken, A. L. Severance, A. Ralston, J. G. Knott, and K. E. Latham. Novel key roles for structural maintenance of chromosome flexible domain containing 1 (Smchd1) during preimplantation mouse development. Molecular Reproductive Development, 85(7):635–648, 2018. ISSN 1098-2795 (Electronic) 1040-452X (Linking). doi: 10.1002/mrd.23001.

38. M. L. Ruebel, K. A. Vincent, P. Z. Schall, K. Wang, and K. E. Latham. SMCHD1 terminates the first embryonic genome activation event in mouse two-cell embryos and contributes to a transcriptionally repressive state. American Journal of Physiology and Cell Physiology, 317(4):C655–C664, 2019. ISSN 1522-1563 (Electronic) 0363-6143 (Linking). doi: 10.1152/ajpcell.00116.2019.

39. C. W. Hanna, R. Perez-Palacios, L. Gahurova, M. Schubert, F. Krueger, L. Biggins, S. Andrews, M. Colome-Tatche, D. Bourc’his, W. Dean, and G. Kelsey. Endogenous retroviral insertions drive non-canonical imprinting in extra-embryonic tissues. Genome Biology, 20(1):225, 2019. ISSN 1474-760X (Electronic) 1474-7596 (Linking). doi: 10.1186/s13059-019-1833-x.

40. T. Nagano, J. A. Mitchell, L. A. Sanz, F. M. Pauler, A. C. Ferguson-Smith, R. Feil, and P. Fraser. The Air noncoding RNA epigenetically silences transcription by targeting G9a to chromatin. Science, 322(5908):1717–20, 2008. ISSN 1095-9203 (Electronic) 0036-8075 (Linking). doi: 10.1126/science.1163802.

41. L. Redrup, M. R. Branco, E. R. Perdeaux, C. Krueger, A. Lewis, F. Santos, T. Nagano, B. S. Cobb, P. Fraser, and W. Reik. The long noncoding RNA Kcnq1ot1 organises a lineage-specific nuclear domain for epigenetic gene silencing. Development, 136(4): 525–30, 2009. ISSN 0950-1991 (Print) 0950-1991 (Linking). doi: 10.1242/dev.031328.

42. F. Sleutels, R. Zwart, and D. P. Barlow. The non-coding Air RNA is required for silencing autosomal imprinted genes. Nature, 415(6873):810–3, 2002. ISSN 0028-0836 (Print) 0028-0836 (Linking). doi: 10.1038/415810a.

43. Daniel Andergassen, Markus Muckenhuber, Philipp C. Bammer, Tomasz M. Kulinski, Hans-Christian Theussl, Takahiko Shimizu, Josef M. Penninger, Florian M. Pauler, and Quanah J. Hudson. The Airn lncrna does not require any DNA elements within its locus to silence distant imprinted genes. PLOS Genetics, 15(7):1–18, 07 2019. doi: 10.1371/journal.pgen.1008268.

44. J. M. Frost and G. E. Moore. The importance of imprinting in the human placenta. PLoS Genetics, 6(7):e1001015, 2010. ISSN 1553-7404 (Electronic) 1553-7390 (Linking). doi: 10.1371/journal.pgen.1001015.

45. Annabelle Lewis, Kohzoh Mitsuya, David Umlauf, Paul Smith, Wendy Dean, Joern Walter, Michael Higgins, Robert Feil, and Wolf Reik. Imprinting on distal chromosome 7 in the placenta involves repressive histone methylation independent of DNA methylation. Nature Genetics, 36(12):1291–1295, 2004. ISSN 1546-1718. doi: 10.1038/ng1468.

46. David Umlauf, Yuji Goto, Ru Cao, Frédérique Cerqueira, Alexandre Wagschal, Yi Zhang, and Robert Feil. Imprinting along the Kcnq1 domain on mouse chromosome 7 involves repressive histone methylation and recruitment of Polycomb group complexes. Nature Genetics, 36(12):1296–1300, 2004. ISSN 1546-1718. doi: 10.1038/ng1467.

47. N. Jansz, T. Nesterova, A. Keniry, M. Iminitoff, P. F. Hickey, G. Pintacuda, O. Masui, S. Kobelke, N. Geoghegan, K. A. Breslin, T. A. Willson, K. Rogers, G. F. Kay, A. H. Fox, H. Koseki, N. Brockdorff, J. M. Murphy, and M. E. Blewitt. Smchd1 targeting to the inactive X is dependent on the Xist-HnrnpK-PRC1 pathway. Cell Reports, 25(7): 1912–1923 e9, 2018. ISSN 2211-1247 (Electronic). doi: 10.1016/j.celrep.2018.10.044.

48. K. U. Wagner, R. J. Wall, L. St-Onge, P. Gruss, A. Wynshaw-Boris, L. Garrett, M. Li, P. A. Furth, and L. Hennighausen. Cre-mediated gene deletion in the mammary gland. Nucleic Acids Research, 25(21):4323–30, 1997. ISSN 0305-1048 (Print) 0305-1048 (Linking). doi: 10.1093/nar/25.21.4323.

49. J. C. de Greef, Y. D. Krom, B. den Hamer, L. Snider, Y. Hiramuki, R. F. P. van den Akker, K. Breslin, M. Pakusch, D. C. F. Salvatori, B. Slutter, R. Tawil, M. E. Blewitt, S. J. Tapscott, and S. M. van der Maarel. Smchd1 haploinsufficiency exacerbates the phenotype of a transgenic FSHD1 mouse model. Human Molecular Genetics, 27(4): 716–731, 2018. ISSN 1460-2083 (Electronic) 0964-6906 (Linking). doi: 10.1093/hmg/ddx437.

50. P. C. Orban, D. Chui, and J. D. Marth. Tissue- and site-specific DNA recombination in transgenic mice. PNAS, 89(15):6861–5, 1992. ISSN 0027-8424 (Print) 0027-8424 (Linking). doi: 10.1073/pnas.89.15.6861.

51. Mark Lewandoski, Karen Montzka Wassarman, and Gail R. Martin. Zp3–cre, a transgenic mouse line for the activation or inactivation of loxp-flanked target genes specifically in the female germ line. Current Biology, 7(2):148–151, 1997. ISSN 09609822. doi: 10.1016/s0960-9822(06)00059-5.

52. H. S. Leong, K. L. Chen, Y. F. Hu, S. Lee, J. Corbin, M. Pakusch, J. M. Murphy, I. J. Majewski, G. K. Smyth, W. S. Alexander, D. J. Hilton, and M. E. Blewitt. Epigenetic regulator Smchd1 functions as a tumor suppressor. Cancer Research, 73(5):1591–1599, 2013. ISSN 0008-5472. doi: 10.1158/0008-5472.Can-12-3019.

53. A. Keniry, L. J. Gearing, N. Jansz, J. Liu, A. Z. Holik, P. F. Hickey, S. A. Kinkel, D. L. Moore, K. Breslin, K. Chen, R. Liu, C. Phillips, M. Pakusch, C. Biben, J. M. Sheridan, B. T. Kile, C. Carmichael, M. E. Ritchie, D. J. Hilton, and M. E. Blewitt. Setdb1-mediated H3K9 methylation is enriched on the inactive X and plays a role in its epigenetic silencing. Epigenetics & Chromatin, 9:16, 2016. ISSN 1756-8935 (Print) 1756-8935 (Linking). doi: 10.1186/s13072-016-0064-6.

54. K. Theiler. The house mouse: Atlas of embryonic development. Springer-Verlag, New York, 1989. ISBN 978-3-642-88418-4.

55. J. Schindelin, I. Arganda-Carreras, E. Frise, V. Kaynig, M. Longair, T. Pietzsch, S. Preibisch, C. Rueden, S. Saalfeld, B. Schmid, J. Y. Tinevez, D. J. White, V. Hartenstein, K. Eliceiri, P. Tomancak, and A. Cardona. Fiji: an open-source platform for biological-image analysis. Nature Methods, 9(7):676–82, 2012. ISSN 1548-7105 (Electronic) 1548-7091 (Linking). doi: 10.1038/nmeth.2019.

56. M. Borensztein, L. Syx, K. Ancelin, P. Diabangouaya, C. Picard, T. Liu, J. B. Liang, I. Vassilev, R. Galupa, N. Servant, E. Barillot, A. Surani, C. J. Chen, and E. Heard. Xist-dependent imprinted X inactivation and the early developmental consequences of its failure. Nature Structural & Molecular Biology, 24(3):226–233, 2017. ISSN 1545-9985 (Electronic) 1545-9985 (Linking). doi: 10.1038/nsmb.3365.

57. Felix Krueger and Simon R. Andrews. SNPsplit: Allele-specific splitting of alignments between genomes with known SNP genotypes. F1000Research, 5, 2016. ISSN 2046-1402. doi: 10.12688/f1000research.9037.1.

58. D. Kim, B. Langmead, and S. L. Salzberg. HISAT: a fast spliced aligner with low memory requirements. Nature Methods, 12(4):357–60, 2015. ISSN 1548-7105 (Electronic) 1548-7091 (Linking). doi: 10.1038/nmeth.3317.

59. R Core Team. R: A Language and Environment for Statistical Computing. R Foundation for Statistical Computing, Vienna, Austria, 2019.

60. Y. Liao, G. K. Smyth, and W. Shi. featurecounts: an efficient general purpose program for assigning sequence reads to genomic features. Bioinformatics, 30(7):923–30, 2014. ISSN 1367-4811 (Electronic) 1367-4803 (Linking). doi: 10.1093/bioinformatics/btt656.

61. Y. Liao, G. K. Smyth, and W. Shi. The R package Rsubread is easier, faster, cheaper and better for alignment and quantification of RNA sequencing reads. Nucleic Acids Research, 47(8):e47, 2019. ISSN 1362-4962 (Electronic) 0305-1048 (Linking). doi: 10.1093/nar/gkz114.

62. Leo. Breiman, Jerome. Friedman, J. Charles. Stone, and R.A. Olshen. Classification and Regression Trees. Wadsworth Statistics/Probability. Taylor & Francis Ltd, 1 edition, 1984. ISBN 9780412048418.

63. Terry Therneau and Beth Atkinson. rpart: Recursive Partitioning and Regression Trees, 2019. R package version 4.1-15.

64. D. J. McCarthy, Y. Chen, and G. K. Smyth. Differential expression analysis of multifactor RNA-Seq experiments with respect to biological variation. Nucleic Acids Research, 40(10):4288–97, 2012. ISSN 1362-4962 (Electronic) 0305-1048 (Linking). doi: 10.1093/nar/gks042.

65. M. D. Robinson, D. J. McCarthy, and G. K. Smyth. edgeR: a Bioconductor package for differential expression analysis of digital gene expression data. Bioinformatics, 26 (1):139–40, 2010. ISSN 1367-4811 (Electronic) 1367-4803 (Linking). doi: 10.1093/bioinformatics/btp616.

66. Mark D. Robinson and Alicia Oshlack. A scaling normalization method for differential expression analysis of RNA-seq data. Genome Biology, 11(3), 2010. ISSN 1465-6906. doi: 10.1186/gb-2010-11-3-r25.

67. A. T. Lun, Y. Chen, and G. K. Smyth. It’s DE-licious: A recipe for differential expression analyses of rna-seq experiments using quasi-likelihood methods in edgeR. Methods Molecular Biology, 1418:391–416, 2016. ISSN 1940-6029 (Electronic) 1064-3745 (Linking). doi: 10.1007/978-1-4939-3578-9_19.

68. Yoav Benjamini and Yosef Hochberg. Controlling the false discovery rate: A practical and powerful approach to multiple testing. Journal of the Royal Statistical Society. Series B (Methodological), 57(1):289–300, 1995. ISSN 00359246.

69. S. Su, C. W. Law, C. Ah-Cann, M. L. Asselin-Labat, M. E. Blewitt, and M. E. Ritchie. Glimma: interactive graphics for gene expression analysis. Bioinformatics, 33(13): 2050–2052, 2017. ISSN 1367-4811 (Electronic) 1367-4803 (Linking). doi: 10.1093/bioinformatics/btx094.

70. Hadley Wickham. ggplot2: Elegant Graphics for Data Analysis. Springer-Verlag New York, 2016. ISBN 978-3-319-24277-4.

71. Claus O. Wilke. cowplot: Streamlined Plot Theme and Plot Annotations for ‘ggplot2’, 2019. R package version 1.0.0.

72. Erik Clarke and Scott Sherrill-Mix. ggbeeswarm: Categorical Scatter (Violin Point) Plots, 2017. R package version 0.6.0.

73. Kamil Slowikowski. ggrepel: Automatically Position Non-Overlapping Text Labels with ‘ggplot2’, 2019. R package version 0.8.1.

74. Viktor Petukhov. ggrastr: Raster layers for ggplot2, 2019. R package version 0.1.7.

75. Raivo Kolde. pheatmap: Pretty Heatmaps, 2019. R package version 1.0.12.

76. Y. Chen, B. Pal, J. E. Visvader, and G. K. Smyth. Differential methylation analysis of reduced representation bisulfite sequencing experiments using edgeR. F1000Res, 6:2055, 2017. ISSN 2046-1402 (Electronic) 2046-1402 (Linking). doi: 10.12688/f1000research.13196.2.

77. Lesnoff, M., Lancelot, and R. aod: Analysis of Overdispersed Data, 2012. R package version 1.3.1.

78. F. Krueger and S. R. Andrews. Bismark: a flexible aligner and methylation caller for Bisulfite-Seq applications. Bioinformatics, 27(11):1571–2, 2011. ISSN 1367-4811 (Electronic) 1367-4803 (Linking). doi: 10.1093/bioinformatics/btr167.

79. S. Andrews. FASTQC. a quality control tool for high throughput sequence data. 2010.

80. B. Langmead and S. L. Salzberg. Fast gapped-read alignment with Bowtie 2. Nature Methods, 9(4):357–9, 2012. ISSN 1548-7105 (Electronic) 1548-7091 (Linking). doi: 10.1038/nmeth.1923.

81. S. Andrews. SeqMonk: a tool to visualise and analyse high throughput mapped sequence. 2007.

82. Iain C. Macaulay, Wilfried Haerty, Parveen Kumar, Yang I. Li, Tim Xiaoming Hu, Mabel J. Teng, Mubeen Goolam, Nathalie Saurat, Paul Coupland, Lesley M. Shirley, Miriam Smith, Niels Van der Aa, Ruby Banerjee, Peter D. Ellis, Michael A. Quail, Harold P. Swerdlow, Magdalena Zernicka-Goetz, Frederick J. Livesey, Chris P. Ponting, and Thierry Voet. G&t-seq: parallel sequencing of single-cell genomes and transcriptomes. Nature Methods, 12:519, 2015. doi: 10.1038/nmeth.3370https://www.nature.com/articles/nmeth.3370#supplementary-information.

83. Christof Angermueller, Stephen J. Clark, Heather J. Lee, Iain C. Macaulay, Mabel J. Teng, Tim Xiaoming Hu, Felix Krueger, Sébastien A. Smallwood, Chris P. Ponting, Thierry Voet, Gavin Kelsey, Oliver Stegle, and Wolf Reik. Parallel single-cell sequencing links transcriptional and epigenetic heterogeneity. Nature Methods, 13:229, 2016. doi: 10.1038/nmeth.3728https://www.nature.com/articles/nmeth. 3728#supplementary-information.

